# A genetic platform for a biocementation bacterium

**DOI:** 10.64898/2025.12.05.692644

**Authors:** Luis Ortiz, Ariela Esmurria, Charlie Gilbert, Alexander Crits-Christoph, Tyler P. Barnum, Christopher P. Mancuso, Shinyoung Clair Kang, Julia Leung, Kathrin Fenn, Melanie B. Abrams, Stephanie L. Brumwell, Henry H. Lee, Nili Ostrov

## Abstract

*Sporosarcina pasteurii* is the most widely studied bacterium for microbially-induced calcium carbonate precipitation (MICP), a process of intense interest for materials and construction applications. Despite two decades of investigation, *S. pasteurii* has remained genetically intractable, limiting our mechanistic understanding of biomineralization pathways and constraining efforts to engineer scalable solutions. Here, we present the first genetic toolkit for *S. pasteurii*, including a stable replicating plasmid, a conjugation-based DNA delivery protocol, engineered inducible promoters, and methods for genome modification. Using homologous recombination, we precisely deleted 5.7 kb of the genome spanning two operons encoding urease activity and demonstrated complete loss of biocementation. We also screened a library of engineered transposon constructs for activity in *S. pasteurii* and generated a genome-wide mutant library with >15,000 unique insertion sites. Using this library, we identified putative genes affecting ureolytic growth, revealing previously inaccessible aspects of *S. pasteurii* genetics. This work establishes *S. pasteurii* as a genetically tractable platform for rational engineering of MICP and constitutes the first genetic modification capability within the *Sporosarcina* genus.

## Introduction

Living organisms assemble structural materials under conditions that often challenge conventional industrial chemistry, such as ambient temperatures, dilute feedstocks, and aqueous environments^1–6^. However, translating these capabilities into practical applications requires both mechanistic understanding and genetic tools to engineer these pathways.

Microbially-induced calcium carbonate precipitation (MICP) is one such process. In MICP, microbes drive the precipitation of calcium carbonate (CaCO₃), binding loose aggregates into solid, stone-like material in ambient conditions^7,8^. This process has attracted commercial interest across diverse applications: sealing leaking oil and gas wells^9^, controlling road dust^10^, stabilizing soils, and producing construction materials^11^. MICP-based biocement is also being explored as a lower-emission alternative to traditional cement, which accounts for 7-8% of global fossil fuel emissions^12,13^.

*Sporosarcina pasteurii*, first isolated in 1889, has become the most widely studied organism for MICP^14–17^. *S. pasteurii* performs MICP via ureolysis: an intracellular multi-protein urease enzyme catalyzes the hydrolysis of urea to ammonia and carbonic acid, and the resulting pH increase drives CaCO₃ formation. Nucleation of calcium carbonate crystals (*e.g.,* calcite) is promoted by calcium ion accumulation on the negatively-charged bacterial cell surface. Because cells and reactants are suspended in solution, CaCO₃ precipitation can occur within small pores and cracks, sealing materials together^18^. *S. pasteurii* is attractive for biocementation due to its tolerance to ammonia and high pH, ability to persist as endospores, lack of pathogenicity, and negatively-charged cell surface^19–21^.

Despite decades of research and commercial interest^22^, no genetic engineering methods have been reported for *S. pasteurii*. This gap limits both mechanistic understanding of biomineralization and engineering of improved strains for industrial deployment. Here, we describe foundational genetic engineering methods for *S. pasteurii* including a DNA delivery protocol, replicating plasmids, constitutive and inducible expression systems, transposon mutagenesis, and targeted genome editing. These tools open new opportunities for dissecting the molecular processes underlying biocement formation and improving strains for industrial uses.

## Results and Discussion

### Identifying a replicating plasmid

We first aimed to identify a functional replicating plasmid in *S. pasteurii*. We used MicrobeMod^23^ software to assess methylation patterns and detected activity of a known Type II restriction-modification (RM) system (**Supplementary Table 1**). This suggested that DNA delivery via conjugation may be preferred in order to circumvent bacterial defense systems^24–27^.

We applied our previously-described POSSUM screen^28,29^ to identify functional origins of replication (ORIs) and selective markers in three *Sporosarcina* strains: two nearly identical *S. pasteurii* strains available from strain banks (ATCC 11859 and DSM 33, both designated as strain ‘Gibson 22’) and a urease-negative strain of the *Sporosarcina* genus classified as *Sporosarcina* sp000813425 that we independently isolated and characterized from a soil sample (CVM220) (**Supplementary Table 2**, **Supplementary Figure 1**). This conjugation-based screen delivers a library of 23 different plasmids from an *E. coli* donor into *S. pasteurii* in a single experiment.

Our assay identified one functional ORI, pTHT15, using spectinomycin (SPEC) selection in *Sporosarcina* CVM220 (plasmid pGL2_356, **Supplementary Figure 2**). The library assay did not yield transconjugants for the two *S. pasteurii* strains using SPEC or any other tested antibiotics. However, we found that a plasmid with the same ORI and an erythromycin (ERM) marker (plasmid pGL2_299, **Supplementary Figure 3**) was functional in both *S. pasteurii* strains when delivered in isolation using an extended liquid selection period. Conjugation to both *S. pasteurii* strains routinely reached ≥10^3^ transconjugants per mL within three days with selection on solid media (**Methods**). We estimated a copy number of 39 (relative to the chromosome) based on whole genome sequencing of an *S. pasteurii* ATCC 11859 transconjugant.

### Design of synthetic inducible promoters for *S. pasteurii*

We next aimed to identify functional promoters for heterologous expression in *S. pasteurii.* We cloned 18 putatively constitutive endogenous *S. pasteurii* promoters upstream of an mScarlet reporter and assessed their strengths using a multiplexed barcoded library assay (**Supplementary Table 3**). Using RNA-seq, we observed promoter strengths spanning three orders of magnitude (**Figure 1A**, **Supplementary Figure 4**, **Supplementary Data File 1**).

**Figure 1.**
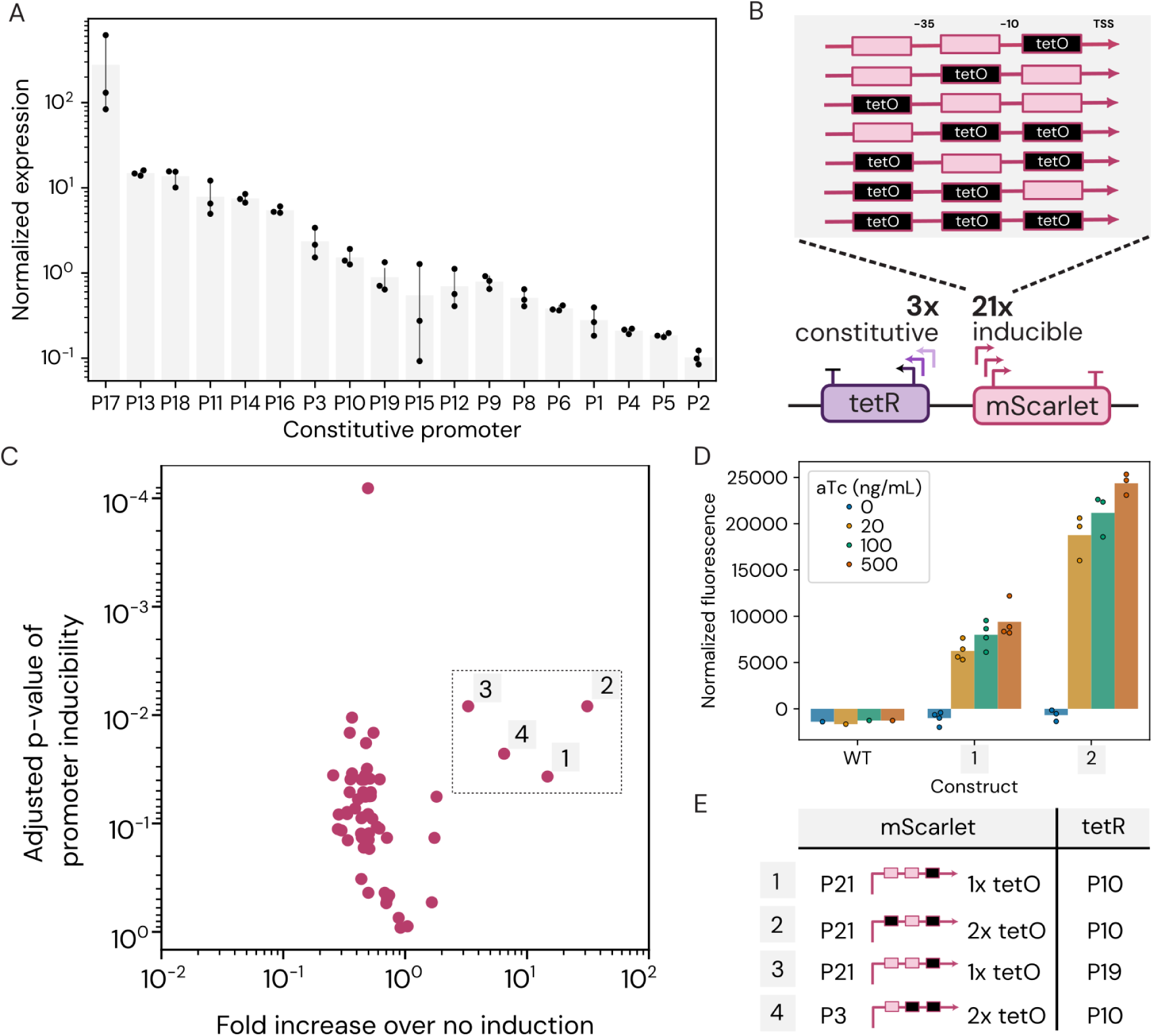
Constitutive and inducible promoter characterization in *S. pasteurii* DSM 33. (**A**) Constitutive promoter expression strength. Rank-ordered DNA-normalized RNA abundance for 18 of 21 constitutive promoter library variants for which sufficient RNA and DNA sequencing data was obtained. Bars represent the mean with 95% confidence intervals (*n*=3). Promoter gene IDs can be seen in Supplementary Table 3. (**B**) Inducible promoter library design. TSS, transcription start site. (**C**) Inducible promoter constructs RNA:DNA sequencing ratio fold change and statistical significance (*n*=3 per condition). The four constructs with significant induction are numbered within the dashed box. (**D**) mScarlet fluorescence of two inducible promoter constructs (1, 2) and wild-type (WT) calculated by dividing background-subtracted red fluorescence (excitation 569 nm, emission 610 nm) by background-subtracted OD_600_ at mid-exponential growth phase (8h) as measured by plate reader. (**E**) Composition of the four inducible promoter constructs identified as functional by sequencing: P21^1tetO^/P10 (1), P21^2tetO^/P10 (2), P21^1tetO^/P19 (3), and P3^2tetO^/P10 (4).

Inspired by previous works^30–32^, we then selected constitutive promoters from this set to engineer inducible promoters. Specifically, we aimed to convert a constitutive promoter to be TetR-repressible, and therefore anhydrotetracycline (aTc)-inducible, by adding tet operator (tetO) sequences. This approach requires a compatible promoter pair: one constitutive promoter to drive TetR expression, and one inducible promoter to drive expression of the mScarlet reporter. From our characterized promoters, we selected three candidates to drive TetR expression: P4 (*rpoB)*, P10 (*GTPase Der*), and P19 (*NADK*) (**Figure 1B**, **Supplementary Table 4**).

To create the inducible promoters, we selected three constitutive promoters - P1 (*EF-Tu*), P3 (*Trigger factor*), and P21 (*tRNA-methionine*) - and designed 21 synthetic variants by inserting one to three tetO sites at different positions relative to the predicted −35 and −10 boxes: upstream, between, or downstream (**Figure 1B**, **Supplementary Table 4**, **Supplementary Data File 2**). We cloned all 63 constitutive-inducible promoter pairs upstream of the mScarlet reporter, with each construct containing a unique nucleic acid barcode for sequencing-based quantification (**Methods**). The pooled library was conjugated into *S. pasteurii* DSM 33, and library sequencing analysis suggested robust conjugative delivery with 98% of all promoter variants represented in the conjugative donor and 93.5% of variants showing up in transconjugants (**Supplementary Figure 5**).

Inducible promoter functionality was evaluated by analyzing the relative abundance of transcript (RNA) and plasmid (DNA) levels with and without aTc. We identified four inducible promoter variants with statistically significant RNA:DNA abundance enrichment when compared to uninduced conditions (t-test; *P*<0.05; FDR=5%), with the best-performing variant (P21^1tetO^/P10) showing a 31.2-fold increase in expression under maximum induction at 20 hours (**Figure 1C**, **Supplementary Figure 6**). Notably, three of these four inducible constructs shared the same TetR promoter (P10) and two shared the same inducible mScarlet promoter (P21^1tetO^). In addition to testing the promoters in a pool, we randomly selected 285 transconjugants and individually screened them on solid media across a range of aTc concentrations. Eight clones exhibited increased fluorescence in the presence of aTc, and sequencing revealed they harbored two of the inducible promoter variants previously identified in the pooled screen: P21^2tetO^/P10 and P21^1tetO^/P10. When assessed in liquid media, both variants showed low fluorescence in the absence of inducer, with the best-performing variant (P21^1tetO^/P10) showing a 41.5-fold increase in expression under maximum induction at 20 hours (**Figure 1D**).

Our results provide a set of 18 constitutive and four inducible promoters with varying expression strengths for heterologous protein expression in *S. pasteurii.* They also highlight the challenges of *de novo* inducible promoter design and the value of a pooled multiplex screening approach for rapid identification of functional promoters.

### Deletion of the urease pathway by homologous recombination

Next, we tested whether we could make genome modifications in *S. pasteurii* using homologous recombination. We delivered a non-replicating plasmid (pGLU_504) carrying an ERM cassette flanked by 1 kb homology sequences targeting the deletion of the urease gene cluster (5703 bp, operons ureABCEF and ureGD) responsible for the ureolytic activity of *S. pasteurii* (**Figure 2A**). The same deletion cassette was used in both *S. pasteurii* strains.

**Figure 2.**
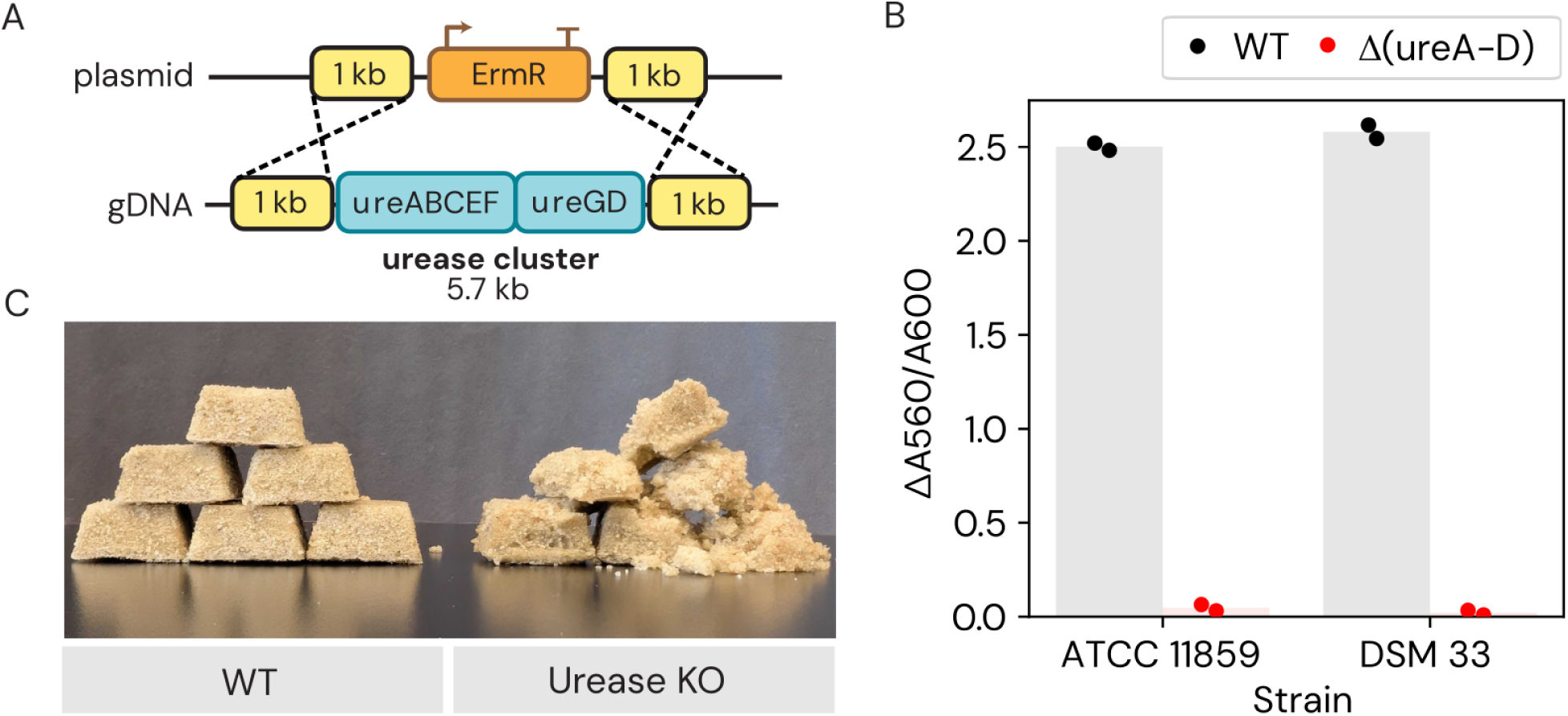
Genomic deletion of the *S. pasteurii* urease pathway. (**A**) Schematic of the urease cluster deletion construct from pGLU_504 plasmid. (**B**) Urease activity of wild-type (WT) and urease-knockout (ΔureA-D) strains as measured by change in phenol red pH indicator (A560 nm / A600 nm) from 0 to 2 hours. (**C**) Biocement-formation assay using *S. pasteurii* DSM 33 wild-type (WT) and urease-knockout (KO) strain.

We isolated 88 ERM-resistant transconjugants for each strain and screened for mutants exhibiting decreased urease activity. Organisms expressing urease can degrade the urea in Stuart’s Urea Broth and produce ammonia that increases the pH from 6.8 to 8.1^33^. We measured the change in pH using phenol red color change (absorbance 560 nm), and found ∼20% of colonies displayed low urease activity (*i.e.,* 19 and 18 colonies in DSM 33 and ATCC 11859, respectively) (**Supplementary Figure 7**, **Methods**). Of these tested colonies, we confirmed by PCR that 50-80% harbored the targeted deletion (*i.e.,* 5/10 and 8/10 colonies in DSM 33 and ATCC 11859, respectively) (**Supplementary Figure 8**). Urease-knockout mutants confirmed by whole genome sequencing lacked any detectable urease activity (**Figure 2B**, **Supplementary Figure 9**).

Finally, we adapted a standard biocement-formation protocol^34^ to test performance of our urease-knockout strains. As expected, wild-type *S. pasteurii* formed adhesive and mechanically stiff biocement. By contrast, urease-knockout *S. pasteurii* exhibited weak adhesion between sand particles, equivalent to negative control samples treated with cementation solution alone (**Figure 2C**). Our assay highlights the essential role of urease in biocement formation and demonstrates the importance of genetic manipulation to study and engineer strains for improved biocementation.

### Construction of a transposon mutant library

Finally, we set out to identify a functional transposon construct for non-targeted insertional mutagenesis using either Himar1 or Tn5. First, we designed an assay to identify functional constructs that effectively drove transposition in *S. pasteurii*. We considered two major common bottlenecks that could impede expression of a transposase in a new bacterial host: (1) codon usage, and (2) promoter compatibility. Given the large combinatorial space of codon usage schemes and promoter sequences that can support a functional transposon, and lack of previously published transposons for *S. pasteurii*, we employed a barcoded pooled library approach to screen transposon variants by sequencing.

For codon adjustment of the transposase gene, we designed and synthesized eight total transposase variants, each with a different codon usage scheme (*i.e.,* five Himar and three Tn5 variants) (**Supplementary Figure 10-11**). Transposase codon usage schemes were generated using optimization to the highest relative synonymous codon usage (RSCU) in *S. pasteurii,* or by codon harmonization^35^ to the relative synonymous codon usage of genomes from which the transposase sequence originated or is highly active in (**Supplementary Table 5**).

We selected 10 endogenous *S. pasteurii* promoters to drive the transposase: eight wild-type promoter sequences and two that were engineered to add RiboJ^36^, a self-cleaving ribozyme that improves mRNA expression (**Supplementary Table 3**, **Supplementary Data File 1**, **Methods**). We cloned all promoters upstream of each codon-adjusted transposase for a total of eight transposase libraries. To select for successful transposition events, we included an ERM marker in the transposon construct. Each of the eight libraries was then conjugated into *S. pasteurii* DSM 33, selected on solid media, and all resulting transconjugants were pooled and sequenced to identify the insertional frequency of each construct.

We observed a range of insertion efficiencies within and between each transposon library (**Figure 3A**). We chose 21 library variants spanning that range (14 Himar and seven Tn5 constructs), and individually conjugated them into *S. pasteurii* DSM 33. We found that the seven constructs that yielded the most colonies had similar insertion efficiency (**Figure 3B**). We selected the top-performing construct (plasmid pGLU_446), a codon-harmonized Himar transposase driven by the RelA/SpoT family protein endogenous promoter (P12), and generated a genome-wide transposon knock-out library in *S. pasteurii* DSM 33 (**Figure 3C**).

**Figure 3.**
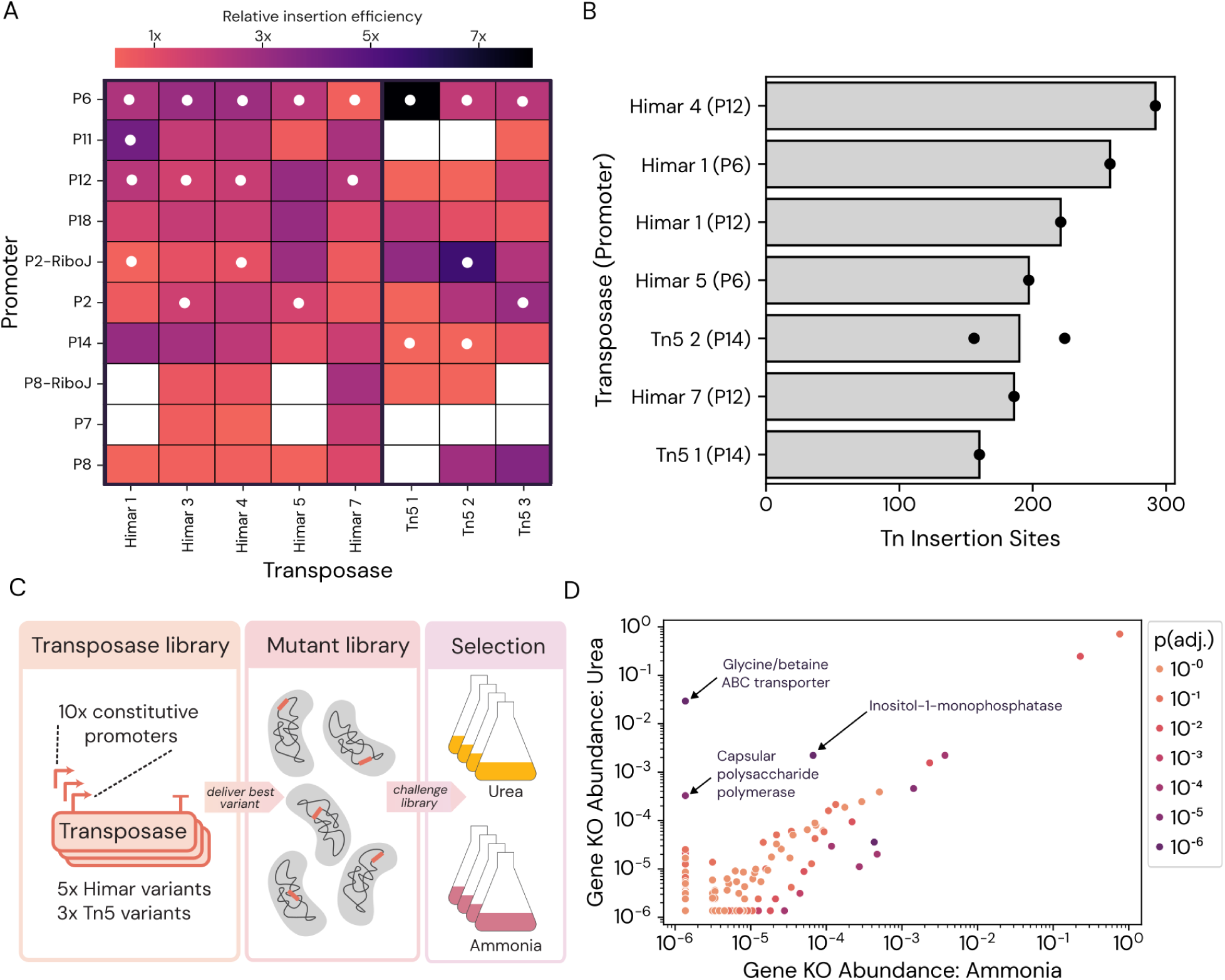
Transposon mutagenesis in *S. pasteurii.* (**A**) Relative insertion efficiency for each of the eight pooled transposase libraries. Each heatmap square represents the frequency of insertions identified with a given promoter-transposase pair relative to the mean number of insertions identified for all promoters with that transposase. White dots signify promoter-transposase constructs that were subsequently delivered in singleton. (**B**) Number of observed unique genomic insertion sites of top-performing promoter-transposase constructs using Tn-seq. (**C**) Overview of transposon library construction and fitness test. (**D**) Tn-seq abundances of transposon library gene mutants after four passages of growth in media containing either urea or ammonia as an abundant nitrogen source.

In our mutant library we identified 21,615 unique insertion sites with low insertional bias across the genome resulting in 92% of genes receiving at least one transposon insertion (**Supplementary Figure 12**). This library included 14,852 unique insertion sites when the library was selected in liquid media, and 8,974 unique insertion sites when the library was selected on solid media. Unexpectedly, we also observed a large proportion of reads mapping to the transposase plasmid sequence (60-90% of total reads), indicating a significant degree of whole-plasmid integration or retention of the transposase plasmid in episomal form. Analysis of selected colonies revealed genomic integration of the plasmid driven by a single-crossover homologous recombination event at the RelA/SpoT family protein endogenous promoter locus in the genome (the same promoter driving the plasmid transposon system). This molecular mechanism likely accounts for the observed plasmid sequencing reads. We thus estimate that 60-90% of this library consists of whole plasmid integration at a single locus, rather than transposon mutants. The extent of this plasmid integration could be reduced in future experiments by using an alternative non-homologous promoter sequence (*e.g.*, a RelA/SpoT family promoter from a close relative of *S. pasteurii*).

We next screened our *S. pasteurii* transposon mutant library in a growth-coupled fitness assay to identify genes impacting *S. pasteurii* ureolysis. We grew the pooled mutant library in a nitrogen-limited media containing either urea or ammonium sulfate as a primary nitrogen source (**Figure 3C-D**). We expected the abundances of mutants that impact ureolysis to be significantly altered in urea compared to an ammonium sulfate control. Four replicate cultures were grown and passaged daily in both urea and ammonia, resulting in ∼7 generations after one day of passage and ∼27 generations after four days of passage. Transposon sequencing was used to identify and quantify the abundances of transposon insertions per gene in each sample.

Overall, we identified eight genes with significant increases in library abundances during growth on urea and five genes with significant increases in library abundances during growth in ammonia (t-test; *P*<0.05; FDR=5%) (**Figure 3D**, **Supplementary Data File 6**). Unexpectedly, we observed two orders of magnitude decrease in library diversity within 24 hours of growth in all conditions and replicates (*i.e.,* decrease in the number of unique insertion sites from 10^4^ to 10^2^) (**Supplementary Figure 13**), likely due to unexpectedly low cell viability post freeze-thaw reducing inoculum population size. Therefore, this screen would be expected to only capture a small subset of variants affecting fitness during ureolysis.

The most significant of the genes identified in our assay was a knockout of gene WP_115361527.1, annotated as a glycine/betaine ABC transporter^37^ of *S. pasteurii* according to the transporter classification database (TCDB)^38^. This transporter class (TCDB 3.A.1.12.1) is highly specific to import of quaternary ammonium compounds^39^, and this gene is genomically adjacent to genes for conversion of choline into glycine betaine. Mutant variants of this gene increased in abundance by over two orders of magnitude after four days of growth with urea, while in ammonia-grown samples, the abundance of these transposon mutants dropped below the limit of detection (**Supplementary Figure 14**). Therefore, knockouts of this gene are expected to exhibit improved growth on urea.

Mutants of a gene encoding inositol-1-monophosphatase (WP_115360339.1) and a gene encoding a capsular polysaccharide polymerase (WP_115359994.1) were also associated with improved growth on urea, albeit less strongly (**Figure 3D**). Gene mutant abundances were fairly similar between the urea and ammonia growth conditions after just one passage, indicating that multiple passages were needed for mutant selection to be detectable (**Supplementary Figure 15**). In sum, this data demonstrates that transposon mutant libraries are a viable method to identify putative gene variants with improved ureolytic growth in *S. pasteurii*. We note that the reduced diversity observed in this initial library screen likely highly limited identification of genes contributing to the ureolytic pathway.

## Conclusions

In this study, we described for the first time a set of fundamental genetic tools for *S. pasteurii*, the most intensely-studied biocement-forming microbe. Our work highlights the value of combinatorial experimental approaches in accelerating discovery in non-model organisms with limited available information. We describe stably replicating plasmids, a DNA delivery method, a set of constitutive and inducible promoters with a large dynamic range, an antibiotic resistance marker, a fluorescent reporter gene, several functional transposon constructs, and targeted genome editing. These resources can be broadly applied to gain deeper insights into the biology underlying biomineralization and generate enhanced, rationally designed biomaterials and construction applications.

## Materials and Methods

### Strains and growth conditions

Unless otherwise noted, *Sporosarcina pasteurii* Gibson 22 (ATCC 11859 and DSM 33) was grown at 30°C in BNH_4_YEM medium, and *Sporosarcina* sp000813425 (CVM220) was grown at 30°C in LB medium. Cultures in flasks were shaken at 225 RPM, and cultures in deepwell plates were shaken at 800 RPM. BNH_4_YEM medium was prepared according to the recipe for ATCC-1376 (“ATCC medium: 1376 *Bacillus pasteurii* NH4-YE medium”). For 1 L of BNH_4_YEM media base, 20 g yeast extract (Fisher Scientific BP9727-2) and 10 g ammonium sulfate (Thermo Scientific AAJ6441936) were either dissolved separately in 500 mL of 0.13 M Tris buffer pH 9, and then autoclaved, or the ammonium sulfate solution was filter sterilized. When preparing Tris buffer with 15.75 g/L Tris Base (Fisher Bioreagents BP152-500), the final media pH would typically reach ∼9, and the pH would not be adjusted; alternatively, Tris-HCl buffer pH 8.5 (Teknova T1085) was diluted to 0.13 M and final media pH was adjusted to ∼9. *Escherichia coli* BW29427 was grown in LB media supplemented with diaminopimelic acid 60 μg/mL (DAP^60^). For antibiotic selection, we use “1X” to refer to the following concentrations: 100 μg/mL kanamycin (KAN), 50 μg/mL spectinomycin (SPEC), 50 μg/mL gentamicin (GEN), 20 μg/mL chloramphenicol (CAM), 20 μg/mL erythromycin (ERM), 50 μg/mL tetracycline (TET); departures from the 1X concentrations are specified in the text.

### Environmental strain isolation

We isolated *Sporosarcina* sp000813425 CVM220 from a rhizosphere soil sample in Watertown, MA, United States. Samples were collected by uprooting the plant, while preserving the root body and attached soil, and documenting photographically for identification via iNaturalist. We diluted 2.88 g of soil in 10 mL of PBS, vortexed for 1 minute to mix with the root body, removed the root body with forceps, and then let the sample settle for 1 minute. Then, we passed 5 mL of the supernatant through a 70 µM cell strainer and incubated at 80°C for 10 minutes using a heat block to kill non-spore formers. We plated on Casein Mannitol (CM) media^40^ (10 g casein digest, 10 g D-mannitol, 5 mg FeSO_4_·7H_2_O, 5 mg ascorbic acid, Trace metal solution (Teknova 1000x Trace Metals Mixture, T1001), 1.5% agarose) to select for relatives of *Bacillus*. On these plates, we spread 100 µL each of 10^-1^ through 10^-4^ dilutions prepared in PBS. We incubated the plates at room temperature for 72 hours, then picked single colonies into a 96-deepwell plate containing 1 mL of CM for growth.

We performed 16S colony PCR to identify the isolates. Each PCR reaction contained 5 µL of DreamTaq 2X Master Mix (Thermo Scientific K1072), 3 µL of molecular grade nuclease-free water (MGW), and 0.5 µL of each of the 10 µM forward and reverse primers (27F and 1492R, (**Supplementary Data File 7**) with 1 µL of cultures diluted 1:10 in MGW as template. The colony PCR protocol was as follows: 95°C for 3 minutes, followed by 30 cycles of a 95°C for 30 seconds, a 30-second anneal at 58°C, and an extension at 72°C for 100 seconds, with a final extension of 72°C for 5 minutes. We sequenced the 16S amplicons and identified the organisms via NCBI BLAST^41^. Through this, we detected an isolate of *Sporosarcina*, which we later confirmed through whole genome sequencing (as described below) as *Sporosarcina* sp00813425 (GTDB).

### DNA handling

#### DNA extraction

Genomic DNA (gDNA) was extracted from *Sporosarcina* strains using the ZYMO Quick-DNA Fungal/Bacterial Miniprep Kit (Zymo Research D6005) or the MagMAX^TM^ Viral/Pathogen Ultra Nucleic Acid Isolation KingFisher kit (Fisher Scientific A42356), according to the manufacturer’s instructions. Nucleic acid concentrations were measured with a Qubit Flex Fluorometer (Fisher Scientific Q45893). Plasmid DNA was extracted from *E. coli* using the QIAprep Spin Miniprep kit (Qiagen 27106).

#### Plasmid cloning

Construction of plasmids via Golden Gate assembly was performed following standard Loop Assembly protocols^28^. Construction of plasmids via isothermal assembly was performed using the NEBuilder HiFi DNA Assembly (New England Biolabs E2621S) according to the manufacturer’s instructions. Plasmid constructs were sequence-verified by performing whole plasmid sequencing (Plasmidsaurus).

### Whole-genome sequencing and analysis

*Sporosarcina* strains (*S. pasteurii* DSM 33, *S. pasteurii* ATCC 11859, *Sporosarcina* sp00813425 CVM220) were sequenced using Oxford Nanopore (R10.4.1) long read sequencing (Plasmidsaurus), and basecalling was performed with Dorado v1. The lowest 5% of reads by quality were removed with Filtlong v0.2.1. Reads were then downsampled to ∼100x coverage, and Flye v2.9.1^42^ was run for assembly, followed by assembly polishing with Medaka v1.8.0(<u>github.com/nanoporetech/medaka</u>) and genome annotation with Bakta v1.6.1^43^. Methylation motifs were called using MicrobeMod run with default settings^23^. Variant calling was performed using breseq v0.39^44^ with default parameters. Whole-genome sequencing revealed that, while near-identical, these strains differ at 42 loci (**Supplementary Data File 5**). Of note were two non-synonymous mutations in the global transcriptional regulator CodY (K160E and K161E) found in *S. pasteurii* DSM 33. This regulator is known to inhibit the transition to stationary phase^45^ in *Bacillus subtilis* and to influence urease gene expression in that species^46^.

### POSSUM screen

We executed the initial high-throughput ORI-marker screen with the POSSUM toolkit as previously described^29^. We screened the plasmid library with the SPEC resistance marker (pGL2_222) for all three *Sporosarcina* strains; for *S. pasteurii* ATCC 11859 and *S. pasteurii* DSM 33, we also screened with the libraries for KAN, GEN, ERM, TET, and CAM resistance markers (plasmid libraries pGL2_217-221). Each antibiotic marker library that was screened contained 23 bacterial origins of replication (ORIs). As a negative control, conjugations were performed with a plasmid with the same antibiotic resistance cassette, but with only the R6K suicide ORI. Briefly, conjugative recipient and donor strains were inoculated into 1 mL cultures (BNH_4_YEM for *S. pasteurii* Gibson 22 strains, LB for *Sporosarcina* CVM220, LB+CAM_20_+DAP_60_ for *E. coli* BW29427 donor strains) and cultured overnight (30°C, 800 RPM shaking in deepwell plates). Donor cultures were pelleted via centrifugation at 4°C and washed in 900 µL PBS; we note that for *Sporosarcina* CVM220 we centrifuged at 3,000 x g on a Hermle Z446-K centrifuge rather than 5,000 x g on an Allegra X-15R. Then, both donor and recipients were pelleted and re-suspended in ∼100 µL of residual supernatant. Donor and recipient re-suspensions were mixed, and then 20 µL was transferred to LB+DAP_60_ agar in a 96-deepwell plate. The plate was sealed with a breathable cloth membrane and incubated for 20 hours at the temperature suited to recipient growth (30°C). Following incubation, 100 µL of PBS was added to each well, and the deepwell plate was shaken for 5 minutes at 800 RPM to resuspend cells from the surface of the agar. An additional 400 µL of PBS was added to the deepwell plate, then 50 µL was inoculated from each culture well to the corresponding position in selective (0.25X, 1X, and 4X antibiotic concentration) and nonselective media. Culture plates were sealed with breathable cloth membrane and incubated at 30°C shaking at 800 RPM for 48-120 hours, with the duration dependent on the time until growth was observed in selective media. OD_600_ was measured at 24 and 48 hours for all experiments, and at one or more of 72, 96, and 120 hours for experiments with longer durations.

The pTHT15 ORI is low-abundance in the POSSUM toolkit (Gilbert et al. 2024), so we reasoned it might be functional in the *S. pasteurii* Gibson 22 strains but below the lower limit of detection in the initial pooled screen. We performed a secondary conjugation screen on *S. pasteurii* ATCC 11859 to identify a functional plasmid. Here, we tested those arrayed members of the POSSUM library (Gilbert et al. 2024) with the pTHT15 ORI: pGL2_246, pGL2_273, pGL2_283, pGL2_299, pGL2_321, and pGL2_356. We used plasmids with a dummy ORI for each antibiotic: pGL2_228, pGL2_175, and pGL2_229-231. After six days, we visually observed a difference in growth between the pGL2_299 and dummy wells, which was subsequently cultured onto an ERM plate. We re-streaked and performed colony PCR (cPCR) on six individual colonies as an initial check for the presence of the plasmid. For this cPCR, we resuspended colonies in 20 µL of MGW, and used 1 µL of that resuspension as a template for PCR with a DreamTaq 2X MasterMix. We amplified a 135 base pair fragment of the plasmid with primers oCG0031 and oCG0037 (**Supplementary Data File 7**), with an initial melt step of 95°C for 10 minutes followed by 30 cycles with a 30 second melt step at 95°C, a 30 second annealing step at 60°C, a 30 second extension at 72°C, and a 1 minute final amplification. As a final check, we performed whole-genome Oxford Nanopore long read sequencing (Plasmidsaurus), which confirmed the transconjugant as *S. pasteurii* and showed the plasmid present and extrachromosomal.

### Conjugation to *S. pasteurii*

Unless specified, conjugations to *S. pasteurii* were performed based on the previous methods^29^, modified as described below. We prepared donor and recipient cultures in liquid media, with a target of at least 5 OD_600_ units (*i.e.*, biomass equivalent to 1 mL of an OD_600_ = 5 culture) of culture per conjugation plate. We harvested cultures by centrifugation at 5,000 x g for 10 minutes at room temperature. We washed pellets once in 5 mL of PBS, then repeated the centrifugation. We resuspended each cell pellet to a final OD_600_ of 100 in one of: nonselective media, PBS, or PBS+DAP_60_. We then mixed 50 µL each of donors and recipients, unless another ratio is indicated. We either mixed directly on the agar of the 10 cm conjugation petri dish or in a sterile 1.5 mL tube prior to plating. We spread the mixed donor and recipient across the surface of the 10 cm conjugation petri dish agar containing DAP_60_, and incubated the conjugation at 30°C for ∼20 hours unless otherwise indicated. After incubation, we scraped the cells from the conjugation plate into 0.75-1.5 mL of PBS. We then plated on selective and nonselective media lacking DAP_60_. We incubated for 3-7 days at 30°C until growth was observed.

We also explored modified conjugation protocols, none of which showed notable improvement. In tests with *S. pasteurii* ATCC 11859 and DSM 33, we achieved similar conjugation yields as on LB+DAP_60_ (≥10^3^ CFU/mL) and BNH_4_YEM+DAP_60_, only if we first applied a heat-shock to *S. pasteurii* and/or adjusted the conjugation media pH to ∼7 rather than pH of ∼9. Without pH-adjustment or heat shock, conjugation yields on BNH_4_YEM+DAP_60_ fell below the limit of detection (10^1^ CFU/mL).

### Endogenous promoter identification

We extracted endogenous promoters from *S. pasteurii* for proteins expected to be expressed constitutively. To find such proteins, we obtained publicly available proteomics measurements for 42 bacteria from 17 phyla^47^ and grouped proteins by their Pfam annotations obtained from UniProt^48^. The protein intensity measurements for each proteome were normalized using a centered-log ratio transform (base 10), then the mean and standard deviation of centered-log ratio intensities across bacteria were calculated for each Pfam. We selected Pfam protein families with a low standard deviation, encoded in approximately single copies (<1.5 proteins in Pfam per genome) and in most of the genomes (>50%), as these Pfams would be specific to a single, common gene and more likely to be constitutively expressed. To extract promoter sequences from these proteins in *S. pasteurii*, we used PyHMMER to annotate proteins with a profile-HMM for each Pfam^49^. We then extracted the sequences upstream (5’) of these genes, requiring a minimum of 100 bp of intergenic sequence. All promoter variants are listed in **Supplementary Data File 1**.

### Promoter library plasmid backbone construction

We first designed a standard destination vector into which all constitutive and inducible promoter variants could be cloned. To construct this destination vector (pGLU_500), we used commercial DNA synthesis (Twist Bioscience) to create a DNA fragment encoding the tetR gene with a downstream terminator but lacking an upstream promoter. In place of this promoter, we added a Golden Gate assembly dropout part, consisting of an sfGFP expression cassette flanked by SapI enzyme restriction sites. This DNA fragment was introduced into plasmid pGL2_299 upstream of the mScarlet gene by NEBuilder HiFi DNA Assembly such that it removed the promoter driving mScarlet expression and arranged the mScarlet and tetR genes in a diverging orientation. SapI restriction sites were designed to allow one-step introduction of a library of bidirectional promoter designs driving expression of tetR and the mScarlet reporter.

### Constitutive and inducible promoter variant library design and construction

To generate a library of inducible promoter designs (*i.e.*, combinations of tetR and mScarlet promoters), we first selected three endogenous *S. pasteurii* promoters to drive tetR expression (P4, P10 and P19) and a fourth random DNA sequence (P0-2) to act as a negative/background control (**Supplementary Data File 2**). Next, we selected a set of three endogenous promoters with discernible conserved promoter elements (−35 and −10 boxes) to act as the basis for aTc-inducible/tetR-repressible promoters (**Supplementary Data File 2**). For each promoter, we defined three possible locations to insert the 19 bp tet operator sequence (tetO1): upstream of the putative −35 box (distal), between the putative −35 and −10 boxes (core), or downstream of the putative −10 box (proximal) (**Supplementary Data File 2**). We then created combinatorial variants of tetR x mScarlet promoter pairs, including versions of each promoter with one, two, or three tetO sequences inserted at all combinations of the three possible locations described above, resulting in a total of 63 candidate inducible promoter designs. Adding relevant controls - such as promoters lacking tetO sites or lacking a tetR promoter entirely - resulted in a final panel of 88 variants (**Supplementary Data File 2**). For constitutive promoter variants, a total of 21 endogenous promoters identified as described above (see “**Endogenous promoter identification**”) were selected for characterization alongside the inducible promoter variants described above. These constitutive promoter variants were designed with no upstream promoter driving tetR expression. This resulted in a final set of 108 promoter library constructs (**Supplementary Data File 3**).

To these promoter constructs, we added unique barcode sequences to enable DNA and RNA quantification by next-generation sequencing. Barcodes were introduced as 15 bp unique sequences within an N-terminal leader peptide added to the mScarlet coding sequence downstream of a 27 bp constant region. Barcode sequences were designed to encode only Thr, Ala, Ser, or Gly amino acids to reduce possible functional effects of variable leader peptides. Next, upstream and downstream sequences were added to introduce SapI cut sites and overhangs compatible with the pGLU_500 destination vector. Finally, constant flanking sequences were added to enable PCR amplification of the entire library. All promoter variants were then synthesized as an oligo pool (Twist Biosciences) (**Supplementary Data File 3**).

Oligo pools were resuspended to 4 nM final concentration in elution buffer (Qiagen 19086), amplified by cycle-limited PCR (18x cycles with Q5 High-Fidelity 2X Master Mix (New England Biolabs M0492L)), purified by gel purification (E-Gel™ CloneWell™ II, Invitrogen G661818), and concentrated using the DNA Clean & Concentrator Kit (Zymo Research D4004). SapI Golden Gate library assemblies and electroporations into the conjugative donor strain *E. coli* BW29427 were performed as described in Gilbert et al. 2024, resulting in ≥700x library coverage. Liquid selection in 50 mL of LB+CAM+DAP_60_ was used to expand the library by growth at 37°C for 16 hours in *E. coli* BW29427 following transformation. A 5 mL aliquot was taken for miniprep and subsequent DNA-seq quantification of the library variant abundance, the remainder of the liquid culture was glycerol-stocked as 1 mL aliquots to be used for subsequent conjugations to *S. pasteurii*.

### Promoter library conjugation to *S. pasteurii*

A single glycerol stock aliquot of the promoter library was thawed and the entire 1 mL used to inoculate 50 mL of LB+CAM+DAP_60_ for 2 hours at 37°C. Conjugations were performed as described under section “**Standard conjugations to *S. pasteurii***” with the only modification being that the entire conjugation mixture was plated across two large (245 mm) square petri dishes for selection. A total population of ≥6000 transconjugants was scraped, pooled, and glycerol-stocked as 1 mL aliquots.

### Promoter library arrayed colony fluorescence assay

A single 1 mL aliquot of the stocked promoter variant library was thawed and plated onto selective media at a range of dilutions to isolate single colonies. After outgrowth, we picked 285 individual colonies, inoculated them into 400 µL of BNH_4_YEM+ERM medium across three 96-well deepwell plates, and allowed them to grow for 3 days at 30°C with shaking at 800 rpm. These cultures were used to inoculate a set of BNH_4_YEM+ERM agar plates with 0, 20, 100, or 500 ng/mL anhydrotetracycline (aTc). Plates were incubated at 30°C and imaged over time using a Phenobooth+ instrument (Singer) under brightfield and red fluorescence imaging settings. Eight individual colonies exhibiting an increase in fluorescence in increasing aTc concentrations were identified. These eight colonies were inoculated into 4 mL BNH_4_YEM+ERM liquid medium alongside a single variant exhibiting high levels of red fluorescence across all conditions as a positive control and a single variant exhibiting no fluorescence across all conditions as a negative control and grown for 4 hours at 30°C. These cultures were back-diluted 1/20 in 4 mL BNH_4_YEM+ERM liquid medium, grown for 5 hours and used to inoculate 200 µL of BNH_4_YEM+ERM liquid medium to a final OD_600_ of 0.05. These cultures were then monitored for OD_600_ and red fluorescence (excitation 569 nm, emission 610 nm) for 48 hours at 30°C in a microplate reader. Finally, mScarlet fluorescence was calculated by dividing background-subtracted red fluorescence by background-subtracted OD_600_ at mid-exponential growth phase. Plasmid DNA was extracted by using the QIAprep Spin Miniprep Kit (QIAGEN 27104) with a 10 min pre-treatment of P1 buffer-resuspended pellets with 1 mg/mL lysozyme (Sigma-Aldrich L6876-1G) and then sequenced to determine the promoter variant identity (Plasmidsaurus).

### Promoter library pooled DNA/RNA-seq assay

For DNA/RNA-seq analysis of promoter expression, a single 1 mL aliquot of the promoter variant library was thawed and used to inoculate triplicate 4 mL liquid cultures of BNH_4_YEM+ERM medium that were grown at 30°C with shaking for 48 hours. Each culture was then used to inoculate 4 mL liquid cultures of BNH_4_YEM+ERM supplemented with 0, 20, 100, and 500 ng/mL aTc to a final OD_600_ of 0.01 and grown for 20 hours at 30°C with shaking. Finally, cultures were pelleted at 4000 x g for 10 minutes and genomic DNA extracted using the ZymoBIOMICS DNA/RNA Miniprep Kit (Zymo Research R2002) with bead beating and RNA extracted using the Quick-RNA Fungal/Bacterial Miniprep Kit (Zymo Research R2014) with DNase I treatment.

For DNA-seq library preparation, the extracted genomic DNA was diluted to 10 ng/µL. For the first PCR reaction, PCR 1 master mix was prepared using the KAPA HiFi HotStart PCR Kit (Roche 07958897001) by mixing 4 µL of 5X KAPA Hifi Fidelity Buffer, 0.6 µL of 10 mM KAPA dNTP, 0.4 µL KAPA Hifi Polymerase (1 U/µL), 1 µL of 20X EvaGreen dye (Biotium 31000), 11.8 µL of MGW, 0.6 µL of oCK.046 Forward Primer (10 µM), and SB0011 Reverse Primer (10 µM) (**Supplementary Data File 7**). 19 µL of the mastermix was added to 1 µL of the diluted gDNA. The PCR amplification was completed in a qTOWER^3^G thermal cycler (Analytik Jena) using the following cycling conditions: initial denaturation at 95 °C for 3 minutes, 30 cycles of 98 °C denaturation for 20 seconds, 67 °C annealing for 15 seconds, and 72 °C extension for 30 seconds, then final extension at 72 °C for 1 minute.

PCR 1 product was diluted 20-fold to be used as template for the second PCR amplification. PCR 2 master mix was prepared in the same way as PCR 1, but without the primers and added to 1 µL of diluted template along with 0.6 µL i5xx (10 µM) and 0.6 µL of i7xx primers (10 µM) (**Supplementary Data File 7**). Amplification was completed in the qTOWER^3^G with the following cycling conditions: initial denaturation at 95°C for 3 minutes, 30 cycles of 98 °C denaturation for 20 seconds, 70 °C annealing for 15 seconds, and 72 °C extension for 30 seconds. Reaction was stopped at 8 cycles when the amplification curve entered the exponential phase.

For RNA-Seq library preparation, reverse transcription on the extracted total RNA was performed to make cDNA using Superscript IV Reverse Transcriptase (Invitrogen 18-090-010) following the official Invitrogen protocol and using oCK.045 for the primer, which included a 10-base unique molecular identifier (UMI) for more accurate quantification of transcripts in downstream analysis. The first PCR was then performed in the same way as PCR 1 in DNA-seq assay with the following modifications: oCK.046 as Forward Primer, i7xx as Reverse Primer, 2 µL cDNA from the reverse transcription reaction as input, 10.8 µL MGW, and an annealing temperature of 66°C. The second PCR was also performed the same as DNA-seq PCR 2 except P7 was used as reverse primer instead of i7xx (**Supplementary Data File 7**).

The barcoded DNA-seq and RNA-seq samples were pooled in equal volume as two separate sets, then purified with DNA Clean & Concentrator-5. The final products were visualized using Invitrogen E-Gel 2% EX agarose gel (Thermo Fisher Scientific G402022). They were then diluted to 4 nM using Resuspension Buffer (Illumina), and the rest of the library preparation steps were completed following the Denature and Dilute Libraries Guide for MiniSeq System by Illumina. The final libraries were sequenced on Illumina MiniSeq with a 300-cycle kit in a 2 x 150 bp paired-end run.

### Promoter library RNA/DNA sequencing analysis

Sequencing reads for both RNA and DNA samples were quality and adapter trimmed using BBDuk from the BBTools package v39.33^50^ with the following settings: k=19, qtrim=r, trimq=10, maq=10, entropy=0.3. Forward and reverse reads were then merged using BBmerge from the BBTools package with default parameters. Promoter barcode sequences were trimmed and extracted using Cutadapt v5.1^51^ with the following parameters: -g atgggttctcaccaccaccaccatcac -a ATGGTGAGTAAGGGTGAAG -M 16 --trimmed-only -n 2. UMIs were extracted from the RNA samples using Cutadapt v5.1^51^ with the following parameters: cutadapt -g ATGGTGAGTAAGGGTGAAG -M 16 -m 8 --trimmed-only -n 1. VSEARCH^52^ was then run to link promoter barcodes to their corresponding promoters using --search_exact. In Python, RNA-seq data was then dereplicated per UMI: only one read was retained for each UMI, therefore removing PCR duplicates from the data. The relative abundance (promoter barcode abundance divided by total number of reads per sample) of each promoter in the RNA was then normalized to the relative abundance of each promoter in the corresponding DNA sample for that same replicate. For each comparison of the induced conditions (n=4 samples per condition) to the uninduced condition, Welch’s t-test (scipy) was run on normalized RNA:DNA promoter abundances to identify promoters significantly enriched in the induced condition. P-values were corrected for multiple hypotheses using the Benjamini-Hochberg Procedure^53^ (FDR=5%) in statsmodels^54^.

### Targeted urease cluster deletion via homologous recombination

Targeted deletion of the *S. pasteurii* urease gene cluster was performed as follows. We isolated gDNA from *S. pasteurii* DSM 33 using the Zymo Quick-DNA Fungal/Bacterial Miniprep Kit. We generated pGLU_504, an R6K suicide plasmid bearing an *oriT* for conjugative delivery, using a single isothermal assembly with 1kb homology arms amplified from upstream and downstream of the ∼6kb urease cluster in the gDNA, and the ERM resistance gene cassette, oriT, and R6K ORI from pGL2_299. We conjugated pGLU_504 into both *S. pasteurii* Gibson 22 strains (DSM 33 and ATCC 11859) and screened 88 ERM-resistant transconjugants per strain for urease activity using a phenol red colorimetric assay. Briefly, single colonies of transconjugants were picked into 500 uL of BNH_4_YEM+ERM5 media in 96-well culture plates (Axygen P-DW-11-C-S) and incubated overnight at 30℃ shaking at 800 RPM. 5 uL of overnight culture was then diluted into 200 uL Stuart’s Urea Broth in a 96-well flat-bottom assay plate, sealed with a clear gas-impermeable seal (Thermo Fisher AB-0558) to prevent ammonia gas exchange between wells, then monitored kinetically for absorbance at 560 and 600 nm, shaking at 30℃ and 800 RPM. A subset of ten transconjugants per strain showing the lowest urease activity were picked and screened by colony PCR using primers oLO.261 and oLO.262 (**Supplementary Data File 7**). Q5 High-Fidelity 2X Master Mix was used for amplification with the following cycling conditions: initial denaturation at 98°C for 30 seconds, 30 cycles of 98°C denaturation for 10 seconds, 60°C annealing for 15 seconds, and 72°C extension for 1 minute, then final extension at 72°C for 10 minutes. Finally, whole genome sequencing (Plasmidsaurus) was performed on two transconjugants to confirm gene deletion.

### Preparing biocement bricks

We inoculated liquid cultures of wild-type *S. pasteurii* and a *S. pasteurii* pGLU_504 transformant (with a knocked-out urease gene cluster) into flasks containing 60 mL of BNH_4_YEM media and cultured at 30°C for 2 days. A volume equivalent to 30 mL of OD_600_ 1 culture was centrifuged at 4000 x g for 10 minutes and resuspended in 30 mL of freshly prepared Cementation Solution ^34^ comprised of 3 g/L nutrient broth (MP Scientific 1007917), 20 g/L urea, 10 g/L ammonium chloride, and 49 g/L calcium chloride dihydrate. Using a repeated pipette, we applied 1.4 mL of resuspended culture, 1.4 mL of cementation solution, or 1.4 mL of water to an arrayed silicone mold containing 4.1 g of washed sea sand (Fisher S25-3) per well. Molds were stored in a chemical safety hood for ammonia to off-gas and for the bricks to dry. We applied 1 mL of freshly prepared cementation solution (or water) to each brick every 24 hours for an additional 5 days. Bricks were allowed to dry for 48 hours before demolding.

### Transposon variant library cloning

We constructed pooled libraries of transposon constructs - with varied transposase and transposase promoter sequences - as follows. First, we built eight destination vectors with the ERM resistance cassette from pGL2_299 as the transposon cargo and no promoter driving the transposase. In place of this promoter, we added a Golden Gate assembly dropout part, consisting of an sfGFP expression cassette flanked by SapI enzyme restriction sites. These SapI sites were designed to allow one-step introduction of a library of barcoded promoters driving expression of the transposase genes. These vectors also contained the oriT for conjugation and *E. coli* propagation elements. Each destination vector had one of eight variants of the Himar or Tn5 coding DNA sequence, with different codon optimization or harmonization (**Supplementary Table 4**). For sequences that were codon-optimized, codons are optimized for maximum availability in *S. pasteurii* according to the indicated organisms’ codon usage pattern (**Supplementary Table 4**). For sequences that were codon-harmonized, the pattern of rare versus common codons in the wild-type transposase coding sequence is mapped using the indicated source organisms’ codon usage pattern, and this pattern of rare versus common codons is then recreated using the genome-wide codon usage pattern of *S. pasteurii* (**Supplementary Table 4**). The wild-type source of the Himar transposase is *Haematobia irritans*; here, we used the codon usage pattern of *Stomoxys calcitrans* in the same subfamily (*Muscinae*). *P. alcaliphila* is an organism in which the Himar transposase was observed to have high activity, and *E. coli* is the source of the Tn5 transposase. Transposase genes were synthesized (Twist Bioscience), and then amplified using Q5 High-Fidelity 2X Master Mix. PCR reactions were run on a 1% E-Gel (Invitrogen A42345) and bands of the correct size were extracted using a QIAquick Gel Extraction Kit (QIAGEN 28704). 0.2 pmol of each purified DNA fragment was added to reactions containing NEBuilder HiFi DNA Assembly Master Mix, mixed well, then incubated at 50°C for 1 hour. Reactions were chilled on ice for 5 minutes, then 1 µL of reaction was added to 10 µL One Shot PIR2 Chemically Competent *E. coli* (Invitrogen C111110) and incubated on ice for 30 minutes. Cells were heat-shocked at 42°C for 10 seconds, then returned to ice for 5 minutes. 100 µL SOC media was then added to the reactions, and tubes were recovered at 30°C for 1 hour. The full volume of each reaction was spread on LB agar plates supplemented with appropriate antibiotics and incubated at 30°C overnight. Individual colonies were used to inoculate 5 mL LB liquid cultures supplemented with antibiotics and grown overnight at 30°C. 100 µL of culture were submitted to whole plasmid sequencing via Plasmidsaurus. Plasmid DNA was purified from correct clones and used for subsequent SapI Type IIS cloning of promoters upstream the transposase genes.

Next, we selected 10 endogenous *S. pasteurii* promoters. Promoters were synthesized as linear fragments (Twist Bioscience), then amplified with primers that incorporate random barcodes to allow Tn-seq identification of the promoter and transposase pair alongside the insertion site. The barcode structure was AATG-[transposase barcode]-GCA-[promoter barcode] where the [transposon barcode] represents a defined 20 base pair identifier for the transposase variant and the [promoter barcode] represents a random 20mer paired to the promoter. Because the barcodes need to be in the transposon itself for Tn-seq analysis, and the two transposases have different mosaic end sequences, we used different primers for Himar and Tn5 to incorporate the random barcodes. For promoter amplification, forward primers oLO.244 (Himar) or oLO.245 (Tn5) were paired with reverse primer oLO.246 (Twist Adapter Rev) (**Supplementary Data File 7**) using Q5 High-Fidelity 2X Master Mix with the following cycling conditions: initial denaturation at 98°C for 30 seconds, 30 cycles of 98°C denaturation for 10 seconds, 60°C annealing for 15 seconds, and 72°C extension for 1 minute, then final extension at 72°C for 10 minutes. PCR amplicons were purified using 1x AMPure XP Beads (Beckman A63881), eluted in TE, quantified using the Qubit™ 1x dsDNA High Sensitivity Kit (Thermo Fisher Scientific Q33231), then pooled equimolar to 50 nM. Promoter pools were cloned via SapI Golden Gate assembly following standard Loop Assembly protocols^28^.

We initially transformed 2 uL of each pooled assembly into 20 uL of a high-copy strain (OneShot PIR1) for our conditional R6K *E. coli* ORI, and selected transformants in 10 mL liquid outgrowths in LB supplemented with 100 μg/mL carbenicillin (CARB). Plasmid DNA was extracted by miniprep using the QIAprep Spin Miniprep Kit (QIAGEN 27104) and sequenced with the Plasmidsaurus Plasmid Sequencing service to characterize the pool composition. Due to a high-number of dropouts, we repeated the transformation, selection, and characterization using a lower-copy strain (OneShot PIR2) for our conditional R6K *E. coli* ORI. This second pool in the PIR2 cloning strain yielded greater diversity of observed constructs.

### Transposon variant screening

We delivered the transposon variant library pools (from PIR1 and PIR2) into both *S. pasteurii* strains (ATCC 11859 and DSM 33) via conjugation and selected transconjugants for each pool on a single agar plate with BNH_4_YEM+ERM5. We then prepared sequencing libraries of both transposons (Himar and Tn5) using MmeI digestions (for Himar transposons) and semi-arbitrary PCR (for Tn5 and Himar transposons); the data quality from the semi-arbitrary PCR was higher than for MmeI in this instance. We assessed the relative efficiency of different constructs using Tn-seq analysis. Constructs for further screening were selected among the best performers identified from plasmid preparations in both *E. coli* cloning strains (PIR1 and PIR2) and both Tn-seq library preparation methods (MmeI and semi-arbitrary PCR). These picks represent constructs corresponding to a range of performance (in terms of relative transposition efficiency) in our highest quality dataset (PIR2 cloning strain, semi-arbitrary PCR library preparation). Plasmid pGLU_446 yielded the highest number of insertions under those conditions (∼500 total, with <5% of reads mapping to plasmid; we note that the rate of reads mapping to plasmid varied considerably over different experiments, even with the same transposase construct).

### Transposon mutant library construction

We generated a transposon mutant library as follows. We delivered plasmid pGLU_446 to *S. pasteurii* DSM 33 via conjugation on 15 LB_DAP60_ agar petri plates. We scraped these plates and selected transconjugants with ERM on both liquid (500 mL, ∼1/10th resuspended conjugation biomass) and solid media (11 BioAssay dishes (Corning 431272), ∼9/10ths of resuspended conjugation biomass). We pooled the selected transconjugants from both liquid and solid outgrowths and sequenced them with MmeI Tn-seq.

### Sequencing library preparation for transposon insertion

gDNA extraction of transposon mutants was performed as noted in the “**DNA extraction**” section above.

For the mariner transposon, we used an mmeI-based library preparation method based on our previously described “Preparation of RB-TnSeq libraries” methods^55^, with one modification to the first PCR amplification step: the JL0102 primer was used instead of the listed JL0003 primer. The annealing temperature of the thermal cycling protocol was accordingly adjusted to 65°C. All other parts of the first PCR amplification and all other steps remained the same as the reference.

For the Tn5 transposon, we used a semi-arbitrary PCR library preparation method. An arbitrary priming reaction was done in a 20 μL volume using the KAPA HiFi HotStart PCR Kit with a recipe slightly modified from the manufacturer’s protocol: 4 μL KAPA HiFi Buffer (5X), 0.6 μL KAPA dNTP Mix (10 mM), 1 μL JL0075 primer (50 μM), 0.8 μL KAPA HiFi HotStart DNA Polymerase (1 U/μL), and the remaining 13.6 μL volume split between the appropriate template DNA volume for 500 ng total input and MGW. The reaction was completed in a qTOWER^3^G thermal cycler using the following thermal protocol: 95°C for 5 minutes and 5 cycles of 98°C denaturation for 20 seconds, 23°C annealing for 30 seconds, 68°C extension for 15 seconds. The entire reaction was then purified through a double-sided SPRIselect (Beckman Coulter B23318) bead clean according to the manufacturer’s protocol using an initial 0.5x bead to sample volume ratio to exclude large fragments above 800 bp, followed by taking the supernatant and adding beads for a 1x final total bead to sample volume ratio to exclude small fragments below 250 bp. The bead cleaned product was resuspended in 12.5 μL MGW.

The eluted DNA was then transferred to a fresh tube for the first PCR amplification step with the KAPA Hifi HotStart PCR Kit and 20X EvaGreen Dye. A 20 µL reaction was prepared according the manufacturer’s protocols using 4 μL KAPA HiFi Buffer (5X), 0.6 μL KAPA dNTP Mix (10 mM), 0.5 μL JL0076 primer (10 μM), 1 μL oCG0036 primer (10 μM), 0.4 μL KAPA HiFi HotStart DNA Polymerase (1 U/μL) and 1 μL EvaGreen Dye (20X). The reaction was completed in a qTOWER^3^G thermal cycler using a standard thermal cycling protocol according to manufacturer’s instructions with a 63°C annealing temperature and 30 second extension time. The progress of each sample was tracked through the amplification curves produced in the qPCR real-time and the reaction was halted upon seeing sure signs of amplification above the reaction’s baseline - typically around 13-16 cycles. The entire reaction was then purified through a double-sided SPRIselect bead clean according to the manufacturer’s protocol using an initial 0.45x bead to sample volume ratio to exclude large fragments above 900 bp, followed by taking the supernatant and adding beads for a 0.9x final total bead to sample volume ratio to exclude small fragments below 275 bp. The bead cleaned product was resuspended in 20 μL MGW.

Next, 1 µL of a 20-fold dilution of the first PCR product as added to the second PCR amplification as template for a standard KAPA HiFi HotStart PCR Kit with EvaGreen Dye qPCR reaction done in a 20 µL volume exactly according to the manufacturer’s protocol using 1 µL 20X EvaGreen dye, 0.6 µL i5xx primer (10 µM), and 0.6 µL JL0102 primer (10 µM). The reaction was completed in a qTOWER^3^G thermal cycler using a standard thermal cycling protocol according to manufacturer’s instructions with a 71°C annealing temperature and 30 second extension time. The reaction was again halted upon seeing sure signs of amplification above the reaction’s baseline - typically around 12 cycles. Finally, 1 µL of a 20-fold dilution of the second PCR product as added to the third PCR amplification as template for a standard KAPA HiFi HotStart PCR Kit with EvaGreen Dye qPCR reaction done in a 20 µL volume exactly according to the manufacturer’s protocol using 1 µL 20X EvaGreen dye, 0.6 µL P5 primer (10 µM), and 0.6 µL i7xx primer (10 µM). The reaction was completed in a qTOWER^3^G thermal cycler using a standard thermal cycling protocol according to manufacturer’s instructions with a 68°C annealing temperature and 30 second extension time. The progress of each sample was tracked through the amplification curves produced in the qPCR real-time and the reaction was halted upon reaching the stage of exponential amplification - typically around an Intensity [I] value of 50000 and generally took about 10 cycles. The entire reaction was then purified through a double-sided SPRIselect bead clean according to the manufacturer’s protocol using an initial 0.4x bead to sample volume ratio to exclude large fragments above 1500 bp, followed by taking the supernatant and adding beads for a 0.8x final total bead to sample volume ratio to exclude small fragments below 300 bp.

DNA libraries were quantified using the Qubit™ 1X dsDNA High Sensitivity Kit. The rest of the library preparation was completed following the Denature and Dilute Libraries Guide for MiniSeq System by Illumina, then loaded and sequenced on an Illumina MiniSeq in a 2 x 150 bp paired-end run.

### Transposon variant Tn-seq analysis

Sequencing reads were quality and adapter trimmed using BBDuk from the BBTools package v39.33^50^ with the following settings: k=19, qtrim=r, trimq=10, maq=10, entropy=0.3. Reads with genomic insertion sequences were identified and trimmed using Cutadapt v5.1^51^ with the parameters: -g AGATGTGTATAAGAGACAG -g tagaccggggacttatcatccaacctgtta --max-ee 10 --trimmed-only. Promoter and transposase barcodes were trimmed using cutadapt with the parameters: -g GGGTCACGCGTAGGACGCGTCGAGTAGGGTAATG --trimmed-only --quiet | cutadapt -a GTTCCGCAGCCACACGCCGTGTGAAGCTGG --trimmed-only --quiet - | cutadapt -g file:tn_barcodes.fasta --trimmed-only. Genomic sequences that were within a Levenshtein distance of 4 from the plasmid sequence were removed to eliminate plasmid sequences while accounting for sequencing error. Only 1 read per unique genomic insertion sequence was retained to quantify the number of unique insertions. The relative abundance of unique insertions per promoter was then calculated for each transposase, and normalized to the mean for that transposase to reflect relative transpositional efficiency with respect to the mean per sample.

### Fitness assay on transposon mutants

We prepared an inoculum containing ∼2 x 10^8^ CFU/mL of the *S. pasteurii* transposition library, as calculated from OD_600_ reading of the thawed glycerol stock, with an equal number of cells from liquid and solid selection. We cultured cells in a modified BNH_4_YEM media containing yeast extract and Tris buffer supplemented with either 50mM ammonium sulfate or 50mM urea. Approximately ∼2 x 10^7^ CFU were inoculated into four replicate 5 mL cultures in ammonium sulfate media and four replicate 5 mL cultures in urea media. Every 24 hours, we measured the OD_600_ of each culture and subcultured ∼10^7^ CFU into 5 mL of fresh media, for a total of ∼25 generations over 4 growth cycles. At each daily timepoint, 1 mL aliquots of each culture were pelleted at 13000 RCF for 2 minutes, washed twice with PBS, and frozen at −80C after removal of PBS supernatant for subsequent DNA extraction and sequencing.

### Tn-seq sequencing analysis

Sequencing reads were quality and adapter trimmed using BBDuk from the BBTools package v39.33^50^ with the following settings: k=19, qtrim=r, trimq=10, maq=10, entropy=0.3. Reads with genomic insertion sequences were identified and trimmed using Cutadapt v5.1^51^ with the settings: --error-rate 0.25, -g tagaccggggacttatcatccaacctgtta, --minimum-length 10, --overlap 4. Genomic insertion sequences were then mapped to the *S. pasteurii* genome using bowtie2 v2.5.4^56^ with default parameters. A custom script with the Pysam library (see **Data and Resource Availability**) filtered mapped reads with a MAPQ score >1 (uniquely mapped positions) and ≥85% identity to the reference genome. Insertions were linked to a gene when they occurred in the middle 80% of a gene, ignoring insertions in the first and last 10% of all basepairs of each gene. Insertion site sequences with a Levenshtein distance of 4 from the plasmid sequence were moved to eliminate plasmid insertion sequences, accounting for sequencing errors. Only insertion sites represented by at least 3 sequencing reads were retained in downstream analyses.

For statistical analysis of the Tn-seq fitness experiment, the relative abundances of insertions by gene were calculated by dividing by the number of reads mapping to insertions linked to a gene by the total number of reads per sample. Zero values were replaced with pseudocounts of the minimum non-zero relative abundance normalized value in the dataset. For each gene with detectable insertions in at least two of the four replicates at passage timepoints 1 and 4, a comparison of the mean abundance of that gene in urea and ammonia samples was performed using the two-sample t-test. P-values were corrected for multiple hypotheses using the Benjamini-Hochberg Procedure^53^ (FDR=5%) in statsmodel. Significant genes were analyzed using PaperBLAST^57^ and SeqHub (https://seqhub.org/) for functional annotation and interpretation.

## Data and Resource Availability

Annotated plasmid sequences from this study are available as GenBank files under ‘Supplementary material’. Plasmids pGL2_299 (replicating plasmid), pGLU_446 (best-performing transposon variant), pGLU_504 (urease knockout plasmid), and pGLU_511_79 and pGLU_511_71 (best-performing inducible promoter plasmids) are being made available through Addgene (see Supplementary Data File 5 for Addgene IDs). Custom code and data files for sequencing analyses are made available at https://github.com/cultivarium/SporosarcinaPasteuriiGenetics.

## Author Contributions

N.O. and H.H.L. conceived of and designed the project. A.E. and K.F. planned and performed experiments identifying a replicating plasmid. M.B.A performed experiments testing conjugation efficiency under standard and adapted conditions. C.G. and T.B. designed promoter library variants. C.G. and C.K. screened and tested promoter library variants. L.O. designed, generated, and tested urease knockouts. C.M. performed biocementation experiments. L.O. designed and performed transposon variant screening and transposon library construction. L.O., A.E., and C.M. performed Tn-seq experiments. C.K. and J.L. performed DNA sequencing workflows. A.C.C. analyzed whole genome sequencing, RNA sequencing, and transposon sequencing data. M.B.A, S.L.B. and all other authors assisted with writing the manuscript.

## Supporting information

Plasmid Sequences

## Acknowledgements

We thank all members of the Cultivarium team for discussions throughout this project. Cultivarium acknowledges support from Eric and Wendy Schmidt as a Convergent Research Focused Research Organization (FRO).

## bioRxiv usage

Cultivarium manuscripts are disseminated as bioRxiv preprints. Comments and feedback are encouraged in the sections provided by bioRxiv.

