## Supplementary Materials for "A genetic platform for a biocementation bacterium"

**Supplementary Figures** **2**
Supplementary Figure 1. Urease activity of Sporosarcina strains as measured by phenol red assay. 2
Supplementary Figure 2. POSSUM screen in three S. pasteurii strains. 2
Supplementary Figure 3. Plasmid map of pGL2_299. 3
Supplementary Figure 4. Expression strength of candidate constitutive promoters in S. pasteurii DSM 33. 3
Supplementary Figure 5. Abundance of promoter library variants in E. coli donor and S. pasteurii. 4
Supplementary Figure 6. Expression strength of four candidate inducible promoters in S. pasteurii DSM 33. 5
Supplementary Figure 7. Urease assay in candidate urease-knockout mutants. 6
Supplementary Figure 8. Colony PCR screening of candidate urease-knockout mutants. 7
Supplementary Figure 9. Urease activity assay applied to wild-type and urease-knockout. 8
Supplementary Figure 10. Sequence alignment of Himar transposase variants. 10
Supplementary Figure 11. Sequence alignment of Tn5 transposase variants. 12
Supplementary Figure 12. S. pasteurii transposon mutant library characteristics. 12
Supplementary Figure 13. Transposon mutant library diversity during screening. 13
Supplementary Figure 14. Transposon mutant frequency of a Glycine/betaine ABC transporter. 13
Supplementary Figure 15. Abundances of transposon variants. 14
**Supplementary Tables** **15**
Supplementary Table 1. MicrobeMod analysis of S. pasteurii restriction-modification systems. 15
Supplementary Table 2. Strains used in this study. 16
Supplementary Table 3. Summary of endogenous promoters used in this study. 17
Supplementary Table 4. Promoter sequences used for design of inducible promoter libraries. 18
Supplementary Table 5. Transposase codon usage variants in transposon libraries. 19
**Supplementary Data Files** **20**
Supplementary Data File 1 - Endogenous Sporosarcina promoters used in this study. 20
Supplementary Data File 2 - Design and sequences of inducible promoter library variants. 20
Supplementary Data File 3 - Details of all inducible and constitutive promoter library sequences included in oligo pool synthesis. 20
Supplementary Data File 4 - Plasmids used in this study. 20
Supplementary Data File 5 - Summary of sequence differences between S. pasteurii strains. 20
Supplementary Data File 6 - Sporosarcina pasteurii Tn-Seq experiment gene abundances. 20
Supplementary Data File 7 - Oligos used in this study. 20
Supplementary Data File 8 - Promoter induction statistics. 20
**Supplementary References** **21**

### Supplementary Figures

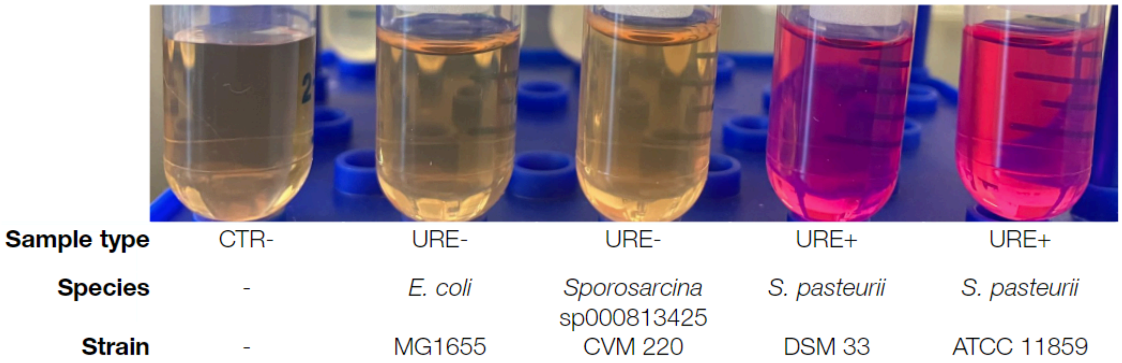


**Supplementary Figure 1. Urease activity of *Sporosarcina* strains as measured by phenol red assay.**
Urease activity of *Sporosarcina* strains was assessed via a phenol red colorimetric assay. In this assay, a change in pH from urease activity results in a color shift from yellow to pink. The positive control *S. pasteurii* Gibson 22 is known to be urease positive (URE+)[1], and the color shift of these strains (ATCC 11859 and DSM 33) is observed as expected. A negative control (CTR-) without any added bacteria resulted in no shift, as expected. A negative control with the naturally urease-deficient (URE-) *E. coli* MG1655 also resulted in no shift, as expected. The soil isolate *Sporosarcina* sp000813425 strain CVM220 also resulted in no shift, indicating no detectable urease activity under the tested conditions.

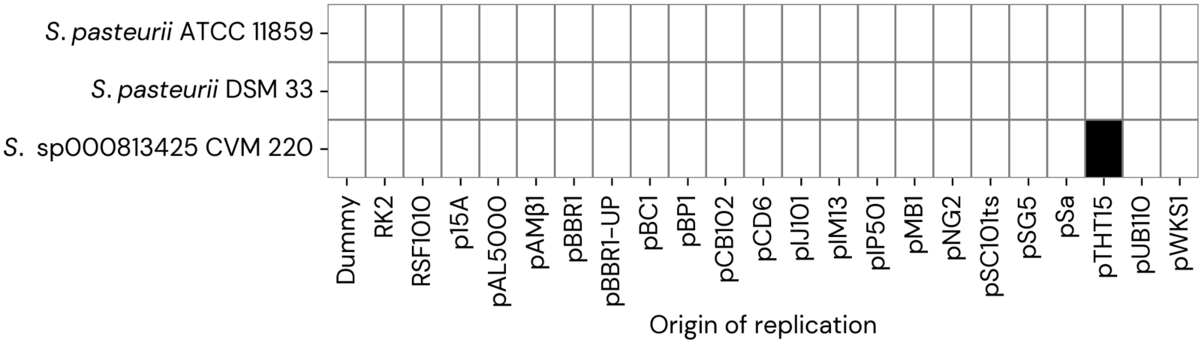


**Supplementary Figure 2. POSSUM screen in three *S. pasteurii* strains.**
Each row corresponds to a strain of *Sporosarcina*, screened with a pooled plasmid library built from the POSSUM toolkit[2], containing 23 different ORIs and a spectinomycin selective marker. Each column indicates an origin of replication within the delivered plasmid libraries. Pooled barcode amplicon sequencing was used to identify successfully replicating plasmids in each strain. An empty box indicates no hit above the limit of detection. A black box indicates a significant hit. One combination, *Sporosarcina* CVM220 with pTHT15, gave a significant hit (plasmid pGL2_356).

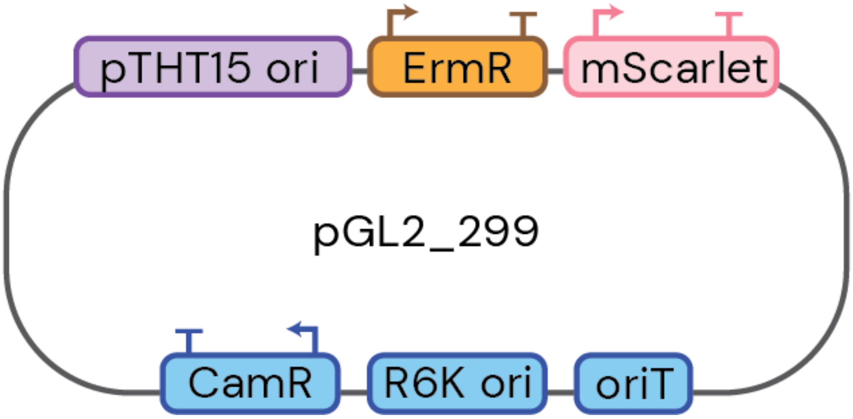


**Supplementary Figure 3. Plasmid map of pGL2_299.**

A plasmid map for pGL2_299, a replicating plasmid functional in both *S. pasteurii* Gibson 22 strains tested (ATCC 11859 and DSM 33). pGL2_299 is a level-two plasmid from our POSSUM toolkit[2]. This plasmid bears a pTHT15 origin of replication, an erythromycin resistance cassette, and an mScarlet fluorescent marker. In addition, a chloramphenicol cassette and R6K origin of replication are included for plasmid maintenance in *E. coli*, and an origin of transfer (*oriT*) is included for conjugative transfer using RP4 (RK2) conjugative machinery in *E. coli*.

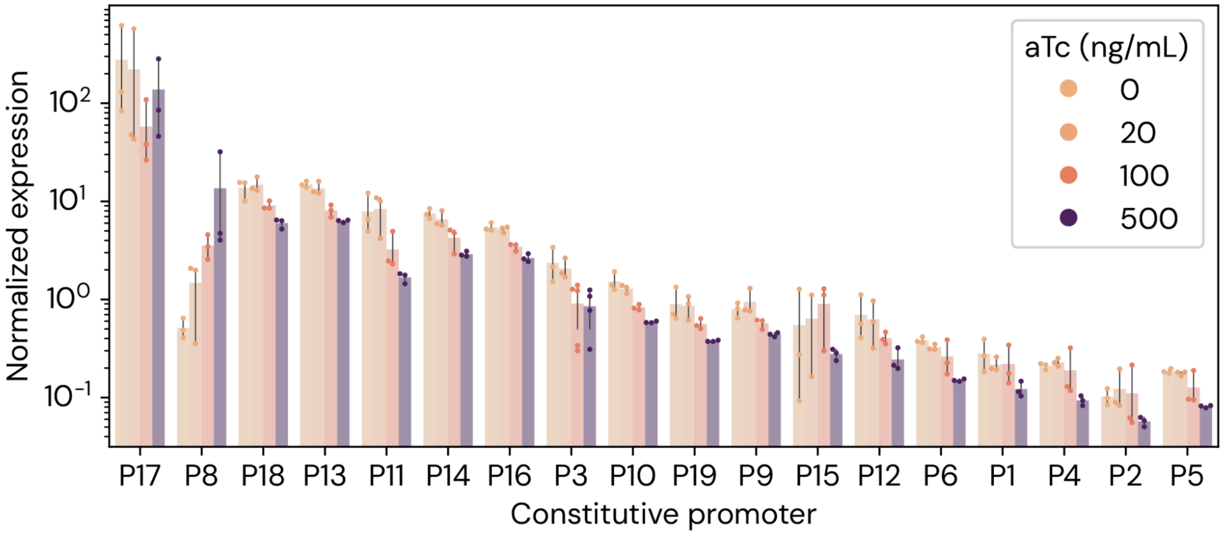


**Supplementary Figure 4. Expression strength of candidate constitutive promoters in *S. pasteurii* DSM 33.**

Normalized expression calculated as DNA-normalized RNA read abundance. Constitutive promoters are arranged in rank order of maximum normalized expression strength. Bars represent the mean values from three replicates and error bars represent 95% confidence intervals. Data was collected in a pooled experiment with candidate inducible promoters, at the concentrations of inducer anhydrotetracycline (aTc) indicated by the bar color.

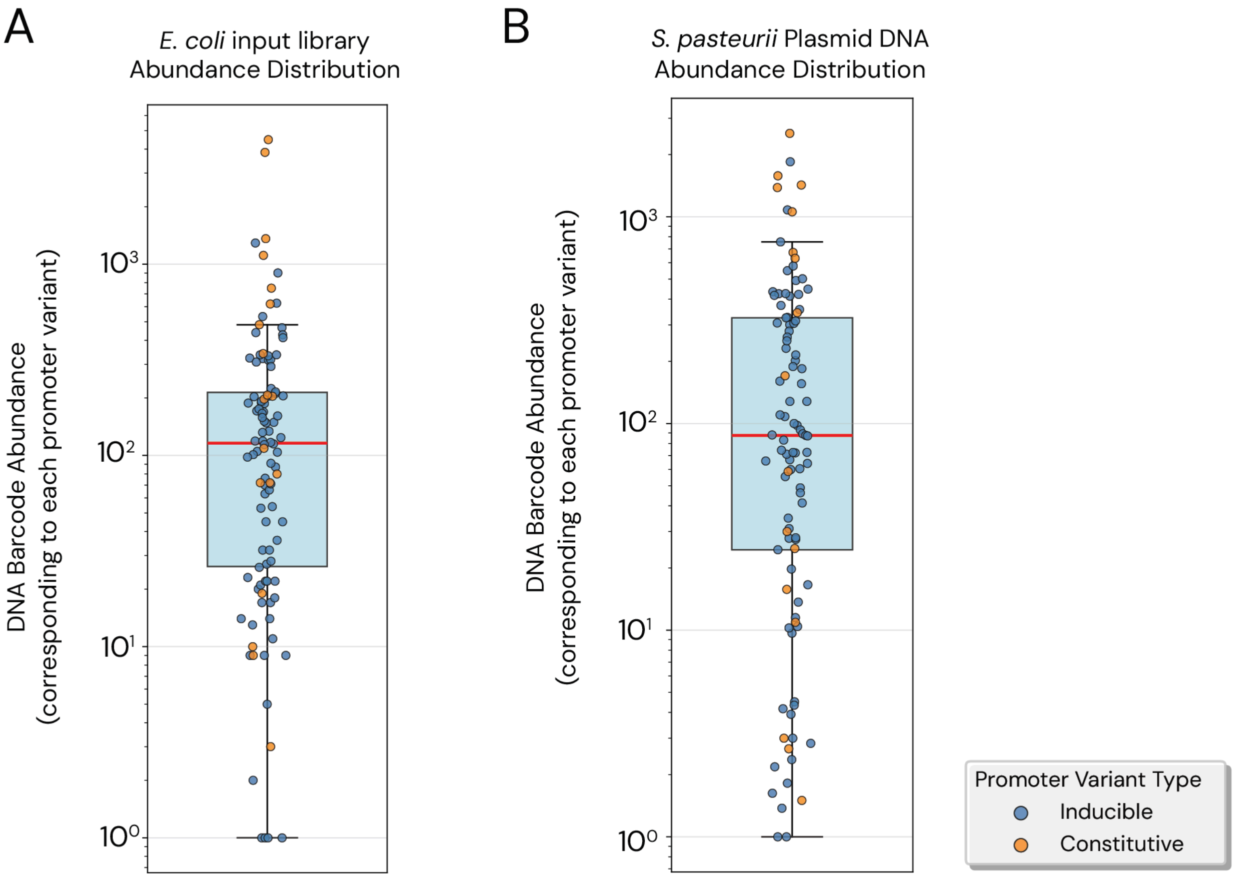


**Supplementary Figure 5. Abundance of promoter library variants in *E. coli* donor and *S. pasteurii.***

DNA barcode abundance for promoter variants in (**A**) the *E. coli* conjugative donor strain and (**B**) *S. pasteurii* DSM 33 following conjugation with *E. coli* harboring the promoter variant library. Barcodes were quantified by amplicon sequencing of the unique barcode region. Data points represent the raw read counts for each promoter variant. *S. pasteurii* samples were prepared as follows: after pooling of promoter library transconjugants, triplicate pooled library cultures were grown, back-diluted, treated with 0, 20, 100, or 500 ng/μL of anhydrotetracycline (aTc), and allowed to grow for a further 20 hours before DNA was extracted for amplicon sequencing. For panel A, two variants with 0 detected reads are not shown, for panel B 10 variants with average abundance <1 are not shown. Box-and-whisker plot shows median value (red line), interquartile range (blue box) and maximum/minimum data points within 1.5×IQR below Q1/Q3 (whiskers).

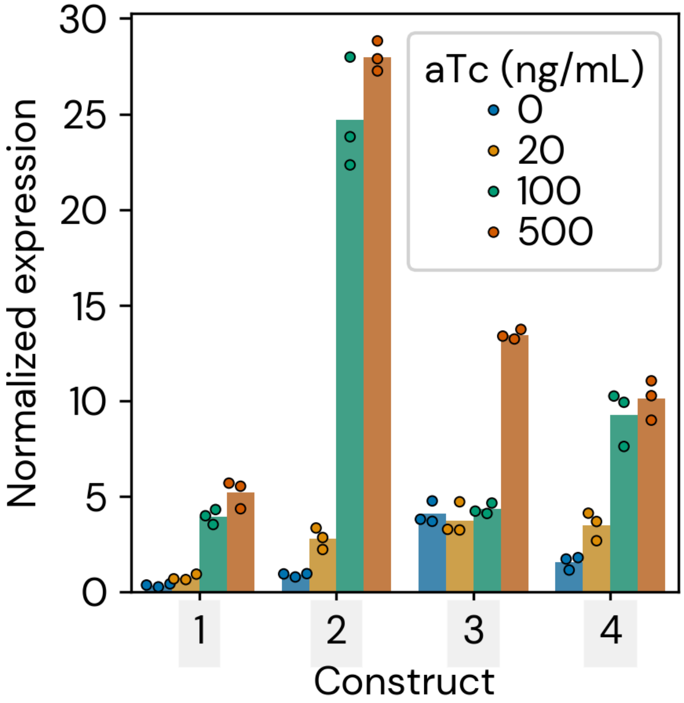


**Supplementary Figure 6. Expression strength of four candidate inducible promoters in *S. pasteurii* DSM 33.**

Normalized expression calculated as DNA-normalized RNA read abundance. The four inducible promoter constructs exhibiting statistically-significant increased expression under maximum induction with anhydrotetracycline (aTc) from the total population are plotted. Construct numbers correspond to **Figure 1C-E**: $P21^{1tetO}$/P10 (1), $P21^{2tetO}$/P10 (2), $P21^{1tetO}$/P19 (3), $P3^{2tetO}$/P10 (4). Bars represent the mean of triplicate measurements.

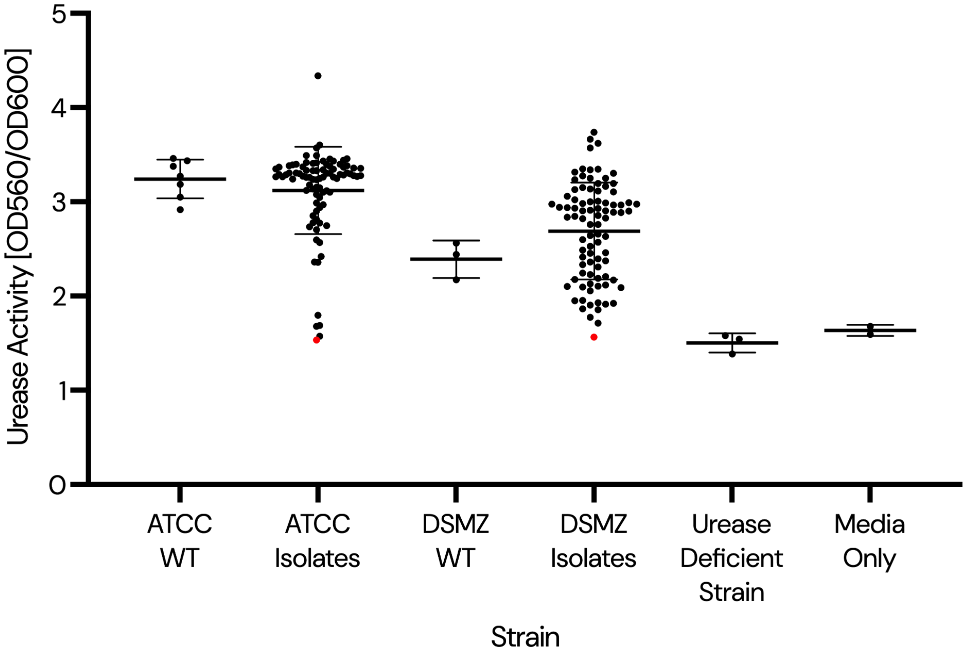


**Supplementary Figure 7. Urease assay in candidate urease-knockout mutants.**
$OD_{600}$-normalized absorbance at $OD_{560}$ for *S. pasteurii* ATCC 11859 and DSM 33 urease-knockout isolates after 2-hour incubation at 30°C. Graph displays mean and standard deviation of samples, with the two isolates ultimately confirmed by whole genome sequencing highlighted in red. The urease-deficient *Sporosarcina* sp000813425 strain CVM220 and media-only controls were used to identify the lower limit of the phenol red colorimetric assay and serve as a threshold to pick putative knockout isolates.

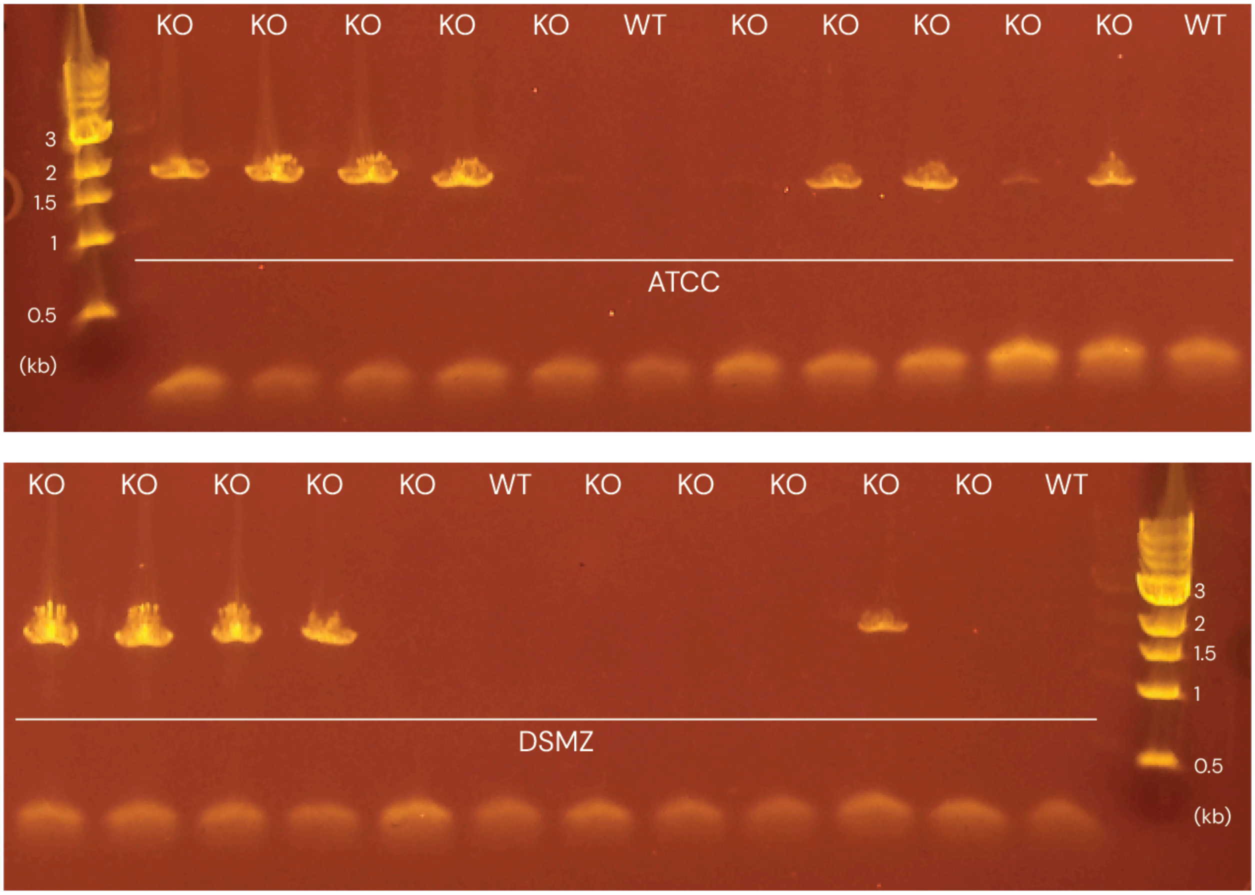


**Supplementary Figure 8. Colony PCR screening of candidate urease-knockout mutants.** PCR results of wild-type (WT) and putative knockout (KO) *S. pasteurii* strains using primers oLO.261 and oLO.261. These primers amplify the junction between genomic DNA and the integrated ERM cassette at the urease gene cluster locus, with on-target integrants yielding a ~2.2 kb band, and WT yielding no band. 8/10 ATCC 11859 isolates, and 5/10 DSM 33 isolates showed the correct band, indicating successful knockout of the urease cluster.

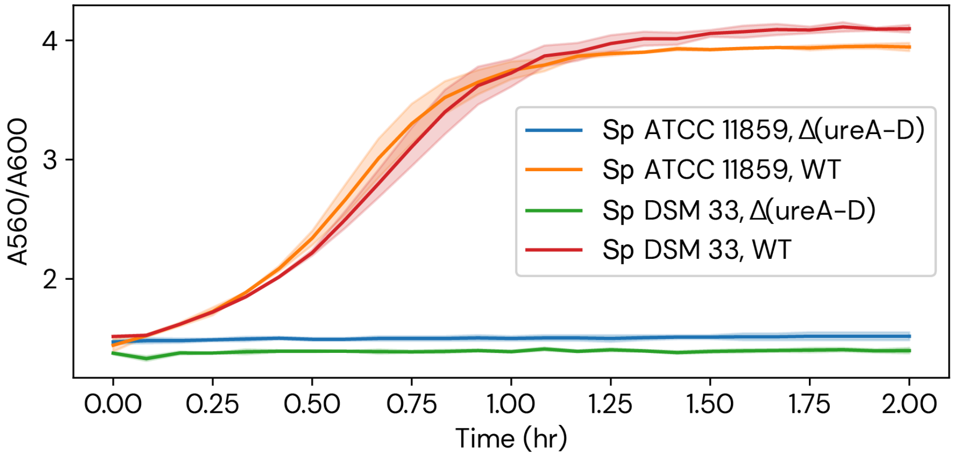


**Supplementary Figure 9. Urease activity assay applied to wild-type and urease-knockout.** Time course urease activity assay (see **Methods**) applied to the wild-type (WT) *S. pasteurii* ('Sp') strains (ATCC 11859 and DSM 33) with and without a knockout of the ~6 kb ureA-D urease genes. Each line represents data from two biological replicates. The x-axis shows time in hours, and the y-axis shows absorbance at 560 nm (proxy for urease activity) normalized to absorbance at 600 nm (proxy for cell density).

```
Himar.001 ATG GAA AAA AAG GAA TTT CGT GTT TTG ATA AAA TAC TGT TTT CTG AAG GGA AAA AAT ACA 60
Himar.003 ATG GAA AAA AAG GAA TTT CGT GTA CTA ATA AAA TAC TGC TTT TTA AAG GGG AAA AAC ACC 60
Himar.004 ATG GAA AAA AAG GAA TTT CGT GTA TTA ATC AAA TAC TGT TTT TTG AAG GGC AAA AAT ACG 60
Himar.005 ATG GAG AAG AAA GAG TTC AGA GTC CTC ATA AAG TAT TGC TTC TTA AAA GGG AAG AAC ACC 60
Himar.007 ATG GAA AAA AAA GAA TTT CGT GTT TTA ATC AAA TAT TGT TTT TTA AAA GGA AAA AAT ACA 60

Himar.001 GTG GAA GCA AAA ACT TGG CTT GAT AAT GAG TTT CCG GAC TCT GCC CCA GGG AAA TCA ACA 120
Himar.003 GTT GAA GCT AAA ACT TGG CTA GAT AAC GAG TTT CCA GAC TCG GCG CCG GGG AAA TCG ACC 120
Himar.004 GTA GAA GCG AAA ACT TGG CTA GAT AAT GAG TTT CCC GAC TCG GCA CCT GGG AAA TCT ACG 120
Himar.005 GTT GAG GCC AAG ACC TGG CTG GAC AAC GAA TTC CCA GAT AGC GCA CCC GGC AAG AGC ACC 120
Himar.007 GTT GAA GCA AAA ACA TGG TTA GAT AAT GAA TTT CCA GAT TCA GCA CCA GGA AAA TCA ACA 120

Himar.001 ATA ATT GAT TGG TAT GCA AAA TTC AAG CGT GGT GAA ATG AGC ACG GAG GAC GGT GAA CGC 180
Himar.003 ATA ATT GAT TGG TAT GCT AAA TTC AAG CGT GGT GAA ATG TCA ACG GAG GAC GGT GAA CGT 180
Himar.004 ATC ATC GAT TGG TAT GCG AAA TTC AAG CGT GGT GAA ATG TCG ACC GAG GAC GGT GAA AGA 180
Himar.005 ATA ATA GAC TGG TAC GCC AAG TTT AAA AGA GGC GAG ATG TCA ACT GAA GAT GGC GAG CGT 180
Himar.007 ATT ATT GAT TGG TAT GCA AAA TTC AAA CGT GGT GAA ATG TCA ACG GAA GAT GGT GAA CGT 180

Himar.001 AGT GGA CGC CCG AAA GAG GTG GTT ACC GAC GAA AAC ATC AAA AAA ATC CAC AAA ATG ATT 240
Himar.003 TCG GGG CGT CCA AAA GAG GTT GTA ACA GAC GAA AAT ATC AAA AAA ATC CAC AAA ATG ATT 240
Himar.004 TCT GGC AGA CCC AAA GAG GTA GTA ACG GAC GAA AAC ATC AAA AAA ATC CAC AAA ATG ATC 240
Himar.005 AGC GGG CGT CCA AAG GAA GTT GTC ACA GAT GAG AAT ATT AAG AAG ATT CAT AAG ATG ATA 240
Himar.007 AGT GGA CGT CCA AAA GAA GTT GTT ACA GAT GAA AAT ATT AAA AAA ATT CAT AAA ATG ATT 240

Himar.001 TTG AAT GAC CGT AAA ATG AAG TTG ATC GAG ATA GCA GAG GCC TTA AAG ATA TCA AAG GAA 300
Himar.003 CTA AAC GAC CGT AAA ATG AAG CTA ATC GAG ATA GCT GAG GCG CTA AAG ATA TCG AAG GAA 300
Himar.004 TTA AAT GAC CGT AAA ATG AAG TTA ATC GAG ATC GCG GAG GCA CTT AAG ATC TCT AAG GAA 300
Himar.005 CTC AAC GAT AGA AAG ATG AAA CTC ATT GAA ATA GCC GAA GCA CTG AAA ATA AGC AAA GAG 300
Himar.007 CTT AAT GAC CGT AAA ATG AAA TTA ATT GAA ATT GCA GAA GCA TTA AAA ATT TCA AAA GAA 300

Himar.001 CGT GTT GGT CAT ATC ATT CAT CAA TAT TTG GAT ATG CGG AAG CTC TGT GCG AAA TGG GTG 360
Himar.003 CGT GTA GGT CAT ATC ATT CAT CAG TAT CTA GAT ATG CGG AAG CTA TGC GCA AAA TGG GTT 360
Himar.004 CGT GTA GGT CAT ATC ATC CAT CAA TAT TTA GAT ATG AGG AAG CTC TGT GCC AAA TGG GTA 360
Himar.005 AGA GTC GGC CAC ATT ATA CAC CAG TAC CTC GAC ATG CGG AAA TTG TGC GCG AAG TGG GTT 360
Himar.007 CGT GTT GGT CAT ATT ATT CAT CAA TAT CTT GAT ATG CGT AAG TTA TGT GCG AAA TGG GTT 360

Himar.001 CCG CGC GAG CTC ACA TTT GAC CAA AAA CAA CGA CGT GTT GAT GAT TCT AAG CGG TGT TTG 420
Himar.003 CCA CGT GAG CTA ACC TTT GAC CAG AAA CAG AGG CGT GTA GAT GAT TCG AAG CGG TGC CTA 420
Himar.004 CCC AGA GAG CTC ACG TTT GAC CAA AAA CAA CGC CGT GTA GAT GAT TCG AAG AGG TGT TTA 420
Himar.005 CCA CGT GAA TTG ACC TTC GAT CAG AAG CAG AGG AGA GTC GAC GAC AGC AAA CGG TGC CTC 420
Himar.007 CCA CGT GAA TTA ACA TTT GAT CAA AAA CAA CGT CGT GTT GAT GAT TCA AAA CGT TGT TTA 420

Himar.001 CAG CTG TTA ACT CGT AAT ACA CCC GAG TTT TTC CGT CGA TAT GTG ACA ATG GAT GAA ACA 480
Himar.003 CAA TTA CTA ACT CGT AAC ACC CCC GAG TTT TTC CGT AGG TAT GTT ACC ATG GAT GAA ACC 480
Himar.004 CAG TTG CTT ACT CGT AAT ACG CCT GAG TTT TTC CGT CGC TAT GTA ACG ATG GAT GAA ACG 480
Himar.005 CAA TTA CTG ACC AGA AAC ACC CCG GAA TTC TTT AGA AGG TAC GTT ACC ATG GAC GAA ACC 480
Himar.007 CAA TTA TTA ACA CGT AAT ACA CCA GAA TTT TTT CGT CGT TAT GTT ACA ATG GAT GAA ACA 480

Himar.001 TGG CTC CAT CAC TAC ACT CCT GAG TCC AAT CGA CAG TCG GCT GAG TGG ACA GCG ACC GGT 540
Himar.003 TGG CTA CAT CAC TAC ACT CCC GAG TCG AAC AGG CAA TCG GCC GAG TGG ACC GCA ACA GGT 540
Himar.004 TGG CTC CAT CAC TAC ACT CCG GAG TCT AAT CGC CAG TCG GCG GAG TGG ACG GCC ACG GGT 540
Himar.005 TGG TTG CAC CAT TAT ACC CCC GAA TCT AAC AGG CAA TCT GCC GAA TGG ACC GCG ACA GGC 540
Himar.007 TGG TTA CAT CAT TAT ACA CCT GAA TCA AAT CGT CAA TCG GCT GAA TGG ACA GCG ACA GGT 540

Himar.001 GAA CCG TCT CCG AAG CGT GGA AAG ACT CAA AAG TCC GCT GGC AAA GTA ATG GCC TCT GTT 600
Himar.003 GAA CCA TCG CCA AAG CGT GGG AAG ACT CAG AAG TCG GCC GGT AAA GTC ATG GCG TCG GTA 600
Himar.004 GAA CCC TCG CCC AAG CGT GGC AAG ACT CAA AAG TCT GCG GGT AAA GTG ATG GCA TCG GTA 600
Himar.005 GAG CCA AGC CCA AAA AGA GGG AAA ACC CAG AAA TCT GCC GGT AAG GTC ATG GCA AGC GTC 600
Himar.007 GAA CCA TCG CCA AAA CGT GGA AAA ACA CAA AAA TCA GCT GGC AAA GTA ATG GCA TCA GTT 600

Himar.001 TTT TGG GAT GCG CAT GGA ATA ATT TTT ATC GAT TAT CTT GAG AAG GGA AAA ACC ATC AAC 660
Himar.003 TTT TGG GAT GCA CAT GGG ATA ATT TTT ATC GAT TAT CTA GAG AAG GGG AAA ACA ATC AAT 660
Himar.004 TTT TGG GAT GCC CAT GGC ATC ATC TTT ATC GAT TAT CTA GAG AAG GGC AAA ACG ATC AAC 660
Himar.005 TTC TGG GAC GCG CAC GGG ATA ATA TTC ATT GAC TAC CTG GAA AAA GGG AAG ACA ATT AAT 660
Himar.007 TTT TGG GAT GCG CAT GGA ATT ATT TTT ATT GAT TAT CTT GAA AAA GGA AAA ACG ATT AAT 660

Himar.001 AGT GAC TAT TAT ATG GCG TTA TTG GAG CGT TTG AAG GTC GAA ATC GCG GCA AAA CGG CCC 720
Himar.003 TCG GAC TAT TAT ATG GCA CTA CTA GAG CGT CTA AAG GTG GAA ATC GCA GCT AAA CGG CCC 720
Himar.004 TCT GAC TAT TAT ATG GCC CTT TTA GAG CGT TTA AAG GTC GAA ATC GCC GCG AAA AGG CCT 720
Himar.005 AGC GAT TAC TAC ATG GCG CTG CTC GAA AGA CTC AAA GTT GAG ATT GCG GCC AAG CGG CCG 720
Himar.007 AGT GAT TAT TAT ATG GCG TTA TTA GAA CGT TTA AAA GTC GAA ATT GCG GCA AAA CGT CCA 720

Himar.001 CAC ATG AAG AAG AAA AAA GTG TTG TTC CAC CAA GAC AAC GCA CCG TGC CAC AAG TCA TTG 780
Himar.003 CAC ATG AAG AAG AAA AAA GTT CTA TTC CAC CAG GAC AAT GCT CCA TGT CAC AAG TCG CTA 780
Himar.004 CAC ATG AAG AAG AAA AAA GTA TTA TTC CAC CAA GAC AAC GCG CCC TGC CAC AAG TCT TTA 780
Himar.005 CAT ATG AAA AAA AAG AAG GTT CTC TTT CAT CAG GAT AAT GCC CCA TGT CAT AAA AGC CTC 780
Himar.007 CAT ATG AAA AAA AAA AAA GTT TTA TTT CAT CAA GAT AAT GCA CCA TGT CAC AAA TCA TTA 780

Himar.001 AGA ACG ATG GCA AAA ATT CAT GAA TTG GGC TTC GAA TTG CTT CCC CAC CCG CCG TAT TCT 840
Himar.003 AGG ACG ATG GCT AAA ATT CAT GAA CTA GGT TTC GAA CTA CTA CCC CAC CCA CCA TAT TCG 840
Himar.004 CGA ACC ATG GCG AAA ATC CAT GAA TTA GGT TTC GAA TTA CTA CCT CAC CCC CCC TAT TCG 840
Himar.005 AGG ACT ATG GCC AAG ATA CAC GAG CTC GGT TTT GAG CTC CTG CCG CAT CCA CCA TAC AGC 840
Himar.007 AGA ACG ATG GCA AAA ATT CAT GAA TTA GGC TTT GAA TTA TTA CCT CAT CCA CCA TAT TCA 840

Himar.001 CCA GAT CTG GCC CCC AGC GAC TTT TTC TTG TTC TCA GAC CTC AAA AGG ATG CTC GCA GGG 900
Himar.003 CCG GAT TTA GCG CCC TCA GAC TTT TTC CTA TTC TCG GAC CTA AAA AGG ATG CTA GCT GGG 900
Himar.004 CCT GAT TTG GCA CCT TCG GAC TTT TTC TTA TTC TCT GAC CTC AAA CGG ATG CTC GCG GGG 900
Himar.005 CCC GAC TTA GCA CCG TCA GAT TTC TTT CTC TTT AGC GAT TTG AAG AGG ATG TTG GCC GGG 900
Himar.007 CCA GAT CTT GCA CCA TCA GAT TTT TTT TTA TTT TCA GAT CTT AAA AGA ATG CTA GCA GGT 900

Himar.001 AAA AAA TTT GGC TGC AAT GAA GAG GTG ATC GCC GAA ACT GAG GCC TAT TTT GAG GCA AAA 960
Himar.003 AAA AAA TTT GGT TGT AAC GAA GAG GTT ATC GCG GAA ACT GAG GCG TAT TTT GAG GCT AAA 960
Himar.004 AAA AAA TTT GGT TGC AAT GAA GAG GTA ATC GCA GAA ACT GAG GCA TAT TTT GAG GCG AAA 960
Himar.005 AAG AAG TTC GGT TGT AAC GAG GAA GTT ATT GCA GAA ACC GAA GCA TAC TTC GAA GCC AAG 960
Himar.007 AAA AAA TTT GGC TGT AAT GAA GAA GTT ATT GCA GAA ACA GAA GCA TAT TTT GAA GCA AAA 960

Himar.001 CCG AAG GAG TAC TAC CAA AAT GGT ATC AAA AAA TTG GAA GGT CGT TAT AAT CGT TGT ATC 1020
Himar.003 CCA AAG GAG TAC TAC CAG AAC GGT ATC AAA AAA CTA GAA GGT CGT TAT AAC CGT TGC ATC 1020
Himar.004 CCC AAG GAG TAC TAC CAA AAT GGT ATC AAA AAA TTA GAA GGT CGT TAT AAT CGT TGT ATC 1020
Himar.005 CCA AAA GAA TAT TAT CAG AAC GGC ATT AAG AAG CTC GAG GGC AGA TAC AAC AGA TGC ATT 1020
Himar.007 CCA AAA GAA TAT TAT CAA AAT GGT ATT AAA AAA TTA GAA GGT CGT TAT AAT CGT TGT ATT 1020

Himar.001 GCT CTT GAA GGG AAC TAT GTT GAA TAA 1047
Himar.003 GCC CTA GAA GGG AAT TAT GTA GAA TAA 1047
Himar.004 GCG CTA GAA GGG AAC TAT GTA GAA TAA 1047
Himar.005 GCC CTG GAG GGG AAT TAC GTC GAG TGA 1047
Himar.007 GCT TTA GAA GGT AAT TAT GTT GAA TAA 1047
```

**Supplementary Figure 10. Sequence alignment of Himar transposase variants.**

Sequences of codon-optimized and codon-harmonized variants of Himar were aligned for visualization. Alignments were performed with the unmodified reference sequences using the Clustal Omega algorithm[3] with mBed-like clustering guide-tree and clustering iteration, 0 combined iterations, and no maximum guide tree iterations or HMM iterations. Positions with modified nucleotides in at least one variant are highlighted in purple.

```
Tn5.001 ATG ATT ACC AGT GCA CTG CAT CGT GCG GCG GAT TGG GCG AAA AGC GTG TTT TCT AGT GCT 60
Tn5.002 ATG ATT ACA TCA GCA TTA CAT CGT GCA GCA GAT TGG GCA AAA TCA GTT TTT TCA TCA GCA 60
Tn5.003 ATG ATT ACA TCG GCT TTA CAT CGT GCA GCA GAT TGG GCA AAA TCA GTT TTT TCG TCG GCC 60

Tn5.001 GCG CTG GGT GAT CCG CGT CGT ACC GCG CGT CTG GTG AAT GTT GCG GCG CAA CTG GCC AAA 120
Tn5.002 GCA TTA GGT GAT CCA CGT CGT ACA GCA CGT TTA GTT AAT GTT GCA GCA CAA TTA GCA AAA 120
Tn5.003 GCA TTA GGT GAT CCA CGT CGT ACA GCA CGT TTA GTT AAC GTA GCA GCA CAG TTA GCG AAA 120

Tn5.001 TAT AGC GGC AAA AGC ATT ACC ATT AGC AGC GAA GGC AGC AAA GCC ATG CAG GAA GGC GCG 180
Tn5.002 TAT TCA GGT AAA TCA ATT ACA ATT TCA TCA GAA GGT TCA AAA GCA ATG CAA GAA GGT GCA 180
Tn5.003 TAT TCA GGT AAA TCA ATT ACA ATT TCA TCA GAA GGT TCA AAA GCG ATG CAA GAA GGT GCA 180

Tn5.001 TAT CGT TTT ATT CGT AAT CCG AAC GTG AGC GCG GAA GCG ATT CGT AAA GCG GGT GCC ATG 240
Tn5.002 TAT CGT TTT ATT CGT AAT CCA AAT GTT TCA GCA GAA GCA ATT CGT AAA GCA GGT GCA ATG 240
Tn5.003 TAT CGT TTT ATT CGT AAC CCA AAT GTT TCA GCA GAA GCA ATT CGT AAA GCA GGT GCG ATG 240

Tn5.001 CAG ACC GTG AAA CTG GCC CAG GAA TTT CCG GAA CTG CTG GCA ATT GAA GAT ACC ACC TCT 300
Tn5.002 CAA ACA GTT AAA TTA GCA CAA GAA TTT CCA GAA TTA TTA GCA ATT GAA GAT ACA ACA TCA 300
Tn5.003 CAA ACA GTT AAA TTA GCG CAA GAA TTT CCA GAA TTA TTA GCT ATT GAA GAT ACA ACA TCG 300

Tn5.001 CTG AGC TAT CGT CAT CAG GTG GCG GAA GAA CTG GGC AAA CTG GGT AGC ATT CAG GAT AAA 360
Tn5.002 TTA TCA TAT CGT CAT CAA GTT GCA GAA GAA TTA GGT AAA TTA GGT TCA ATT CAA GAT AAA 360
Tn5.003 TTA TCA TAT CGT CAT CAA GTT GCA GAA GAA TTA GGT AAA TTA GGT TCA ATT CAA GAT AAA 360

Tn5.001 AGC CGT GGT TGG TGG GTG CAT AGC GTG CTG CTG CTG GAA GCG ACC ACC TTT CGT ACC GTG 420
Tn5.002 TCA CGT GGT TGG TGG GTT CAT TCA GTT TTA TTA TTA GAA GCA ACA ACA TTT CGT ACA GTT 420
Tn5.003 TCA CGT GGT TGG TGG GTT CAT TCA GTT TTA TTA TTA GAA GCA ACA ACA TTT CGT ACA GTT 420

Tn5.001 GGC CTG CTG CAT CAA GAA TGG TGG ATG CGT CCG GAT GAT CCG GCG GAT GCG GAT GAA AAA 480
Tn5.002 GGT TTA TTA CAT CAA GAA TGG TGG ATG CGT CCA GAT GAT CCA GCA GAT GCA GAT GAA AAA 480
Tn5.003 GGT TTA TTA CAT CAG GAA TGG TGG ATG CGT CCA GAT GAT CCA GCA GAT GCA GAT GAA AAA 480

Tn5.001 GAA AGC GGC AAA TGG CTG GCC GCT GCT GCA ACT TCG CGT CTG AGA ATG GGC AGC ATG ATG 540
Tn5.002 GAA TCA GGT AAA TGG TTA GCA GCA GCA GCA ACA TCA CGT TTA CGT ATG GGT TCA ATG ATG 540
Tn5.003 GAA TCA GGT AAA TGG TTA GCG GCC GCC GCT ACT TCG CGT TTA AGG ATG GGT TCA ATG ATG 540

Tn5.001 AGC AAC GTG ATT GCG GTG TGC GAT CGT GAA GCG GAT ATT CAT GCG TAT CTG CAA GAT AAA 600
Tn5.002 TCA AAT GTT ATT GCA GTT TGT GAT CGT GAA GCA GAT ATT CAT GCA TAT TTA CAA GAT AAA 600
Tn5.003 TCA AAT GTT ATT GCA GTT TGT GAT CGT GAA GCA GAT ATT CAT GCA TAT TTA CAG GAT AAA 600

Tn5.001 CTG GCC CAT AAC GAA CGT TTT GTG GTG CGT AGC AAA CAT CCG CGT AAA GAT GTG GAA AGC 660
Tn5.002 TTA GCA CAT AAT GAA CGT TTT GTT GTT CGT TCA AAA CAT CCA CGT AAA GAT GTT GAA TCA 660
Tn5.003 TTA GCG CAT AAT GAA CGT TTT GTT GTT CGT TCA AAA CAT CCA CGT AAA GAT GTT GAA TCA 660

Tn5.001 GGC CTG TAT CTG TAT GAT CAC CTG AAA AAC CAG CCG GAA CTG GGC GGC TAT CAG ATT AGC 720
Tn5.002 GGT TTA TAT TTA TAT GAT CAT TTA AAA AAT CAA CCA GAA TTA GGT GGT TAT CAA ATT TCA 720
Tn5.003 GGT TTA TAT TTA TAT GAT CAC TTA AAA AAT CAA CCA GAA TTA GGT GGT TAT CAA ATT TCA 720

Tn5.001 ATT CCG CAG AAA GGC GTG GTG GAT AAA CGT GGC AAA CGT AAA AAC CGT CCG GCG CGT AAA 780
Tn5.002 ATT CCA CAA AAA GGT GTT GTT GAT AAA CGT GGT AAA CGT AAA AAT CGT CCA GCA CGT AAA 780
Tn5.003 ATT CCA CAA AAA GGT GTT GTT GAT AAA CGT GGT AAA CGT AAA AAT CGT CCA GCA CGT AAA 780

Tn5.001 GCG AGC CTG AGC CTG CGT AGC GGC CGT ATT ACC CTG AAA CAG GGC AAC ATT ACC CTG AAC 840
Tn5.002 GCA TCA TTA TCA TTA CGT TCA GGT CGT ATT ACA TTA AAA CAA GGT AAT ATT ACA TTA AAT 840
Tn5.003 GCA TCA TTA TCA TTA CGT TCA GGT CGT ATT ACA TTA AAA CAA GGT AAT ATT ACA TTA AAT 840

Tn5.001 GCG GTG CTG GCC GAA GAA ATT AAT CCG CCG AAA GGC GAA ACC CCG CTG AAA TGG CTG CTG 900
Tn5.002 GCA GTT TTA GCA GAA GAA ATT AAT CCA CCA AAA GGT GAA ACA CCA TTA AAA TGG TTA TTA 900
Tn5.003 GCA GTT TTA GCG GAA GAA ATT AAC CCA CCA AAA GGT GAA ACA CCA TTA AAA TGG TTA TTA 900

Tn5.001 CTG ACC AGC GAG CCG GTG GAA AGT CTG GCC CAA GCG CTG CGT GTG ATT GAT ATT TAT ACC 960
Tn5.002 TTA ACA TCA GAA CCA GTT GAA TCA TTA GCA CAA GCA TTA CGT GTT ATT GAT ATT TAT ACA 960
Tn5.003 TTA ACA TCA GAG CCA GTT GAA TCG TTA GCG CAG GCA TTA CGT GTT ATT GAT ATT TAT ACA 960

Tn5.001 CAT CGT TGG CGC ATT GAA GAA TTT CAC AAA GCG TGG AAA ACG GGT GCG GGT GCG GAA CGT 1020
Tn5.002 CAT CGT TGG CGT ATT GAA GAA TTT CAT AAA GCA TGG AAA ACA GGT GCA GGT GCA GAA CGT 1020
Tn5.003 CAT CGT TGG CGT ATT GAA GAA TTT CAC AAA GCA TGG AAA ACG GGT GCA GGT GCA GAA CGT 1020

Tn5.001 CAG CGT ATG GAA GAA CCG GAT AAC CTG GAA CGT ATG GTG AGC ATT CTG AGC TTT GTG GCG 1080
Tn5.002 CAA CGT ATG GAA GAA CCA GAT AAT TTA GAA CGT ATG GTT TCA ATT TTA TCA TTT GTT GCA 1080
Tn5.003 CAA CGT ATG GAA GAA CCA GAT AAT TTA GAA CGT ATG GTT TCA ATT TTA TCA TTT GTT GCA 1080

Tn5.001 GTG CGT CTG CTG CAA CTG CGT GAA TCT TTT ACT CCG CCG CAA GCA CTG CGT GCG CAG GGC 1140
Tn5.002 GTT CGT TTA TTA CAA TTA CGT GAA TCA TTT ACA CCA CCA CAA GCA TTA CGT GCA CAA GGT 1140
Tn5.003 GTT CGT TTA TTA CAG TTA CGT GAA TCG TTT ACT CCA CCA CAG GCT TTA CGT GCA CAA GGT 1140

Tn5.001 CTG CTG AAA GAA GCG GAA CAC GTT GAA AGC CAG AGC GCG GAA ACC GTG CTG ACC CCG GAT 1200
Tn5.002 TTA TTA AAA GAA GCA GAA CAT GTT GAA TCA CAA TCA GCA GAA ACA GTT TTA ACA CCA GAT 1200
Tn5.003 TTA TTA AAA GAA GCA GAA CAC GTA GAA TCA CAA TCA GCA GAA ACA GTT TTA ACA CCA GAT 1200

Tn5.001 GAA TGC CAA CTG CTG GGC TAT CTG GAT AAA GGC AAA CGC AAA CGC AAA GAA AAA GCG GGC 1260
Tn5.002 GAA TGT CAA TTA TTA GGT TAT TTA GAT AAA GGT AAA CGT AAA CGT AAA GAA AAA GCA GGT 1260
Tn5.003 GAA TGT CAG TTA TTA GGT TAT TTA GAT AAA GGT AAA CGT AAA CGT AAA GAA AAA GCA GGT 1260

Tn5.001 AGC CTG CAA TGG GCG TAT ATG GCG ATT GCG CGT CTG GGC GGC TTT ATG GAT AGC AAA CGT 1320
Tn5.002 TCA TTA CAA TGG GCA TAT ATG GCA ATT GCA CGT TTA GGT GGT TTT ATG GAT TCA AAA CGT 1320
Tn5.003 TCA TTA CAG TGG GCA TAT ATG GCA ATT GCA CGT TTA GGT GGT TTT ATG GAT TCA AAA CGT 1320

Tn5.001 ACC GGC ATT GCG AGC TGG GGT GCG CTG TGG GAA GGT TGG GAA GCG CTG CAA AGC AAA CTG 1380
Tn5.002 ACA GGT ATT GCA TCA TGG GGT GCA TTA TGG GAA GGT TGG GAA GCA TTA CAA TCA AAA TTA 1380
Tn5.003 ACA GGT ATT GCA TCA TGG GGT GCA TTA TGG GAA GGT TGG GAA GCA TTA CAG TCA AAA TTA 1380

Tn5.001 GAT GGC TTT CTG GCC GCG AAA GAC CTG ATG GCG CAG GGC ATT AAA ATC TAA 1431
Tn5.002 GAT GGT TTT TTA GCA GCA AAA GAT TTA ATG GCA CAA GGT ATT AAA ATT TAA 1431
Tn5.003 GAT GGT TTT TTA GCG GCA AAA GAC TTA ATG GCA CAA GGT ATT AAA ATC TAA 1431
```

**Supplementary Figure 11. Sequence alignment of Tn5 transposase variants.**

Sequences of codon-optimized and codon-harmonized variants of Tn5 were aligned for visualization. Alignments were performed with the unmodified reference sequences using the Clustal Omega algorithm[3] with mBed-like clustering guide-tree and clustering iteration, 0 combined iterations, and no maximum guide tree iterations or HMM iterations. Positions with modified nucleotides in at least one variant are highlighted in purple.

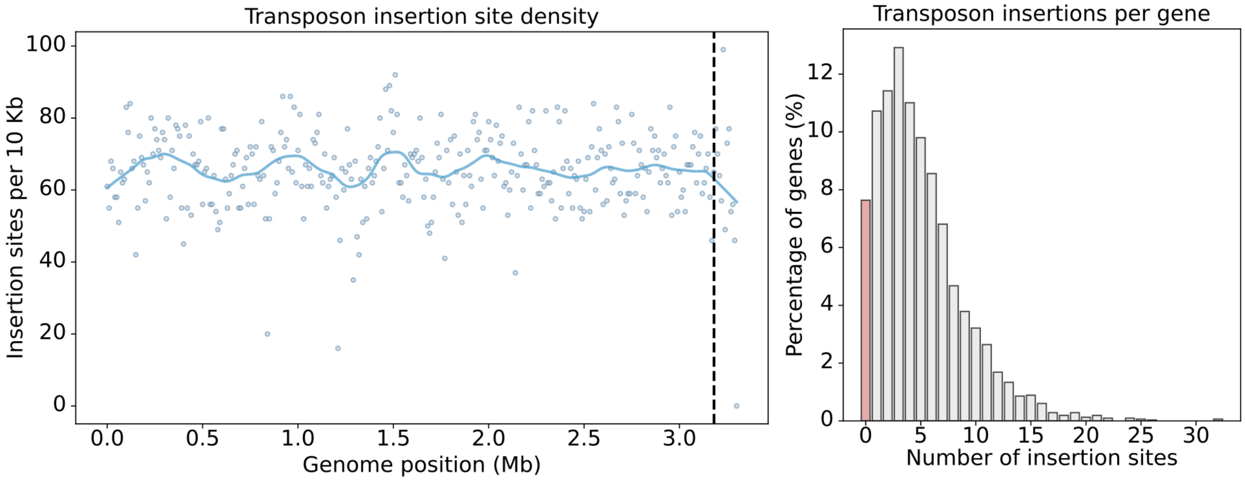


**Supplementary Figure 12. *S. pasteurii* transposon mutant library characteristics.**

Left: The number of unique genomic insertions identified in the library per 10 kb window across the genome. The distinction between the two contigs in our assembly of the genome (which represent the singular *S. pasteurii* DSM 33 chromosome) is visualized with a dividing line. Right: The number of transposon insertions per gene. Only insertions in the middle 80% of a gene are counted. Genes with 0 transposon insertions are marked with the red bar.

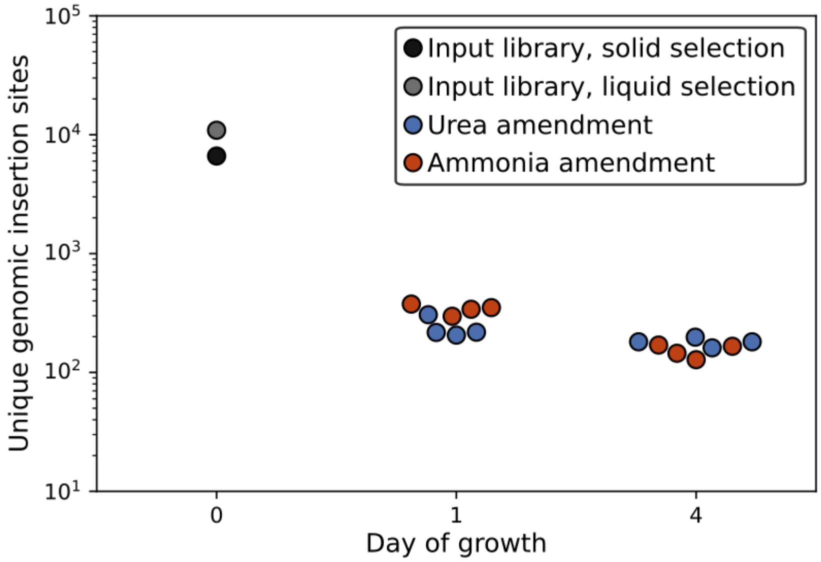


**Supplementary Figure 13. Transposon mutant library diversity during screening.**
The number of unique genomic insertion sites in *S. pasteurii* DSM 33 supported by at least three sequencing reads is shown for each sample.

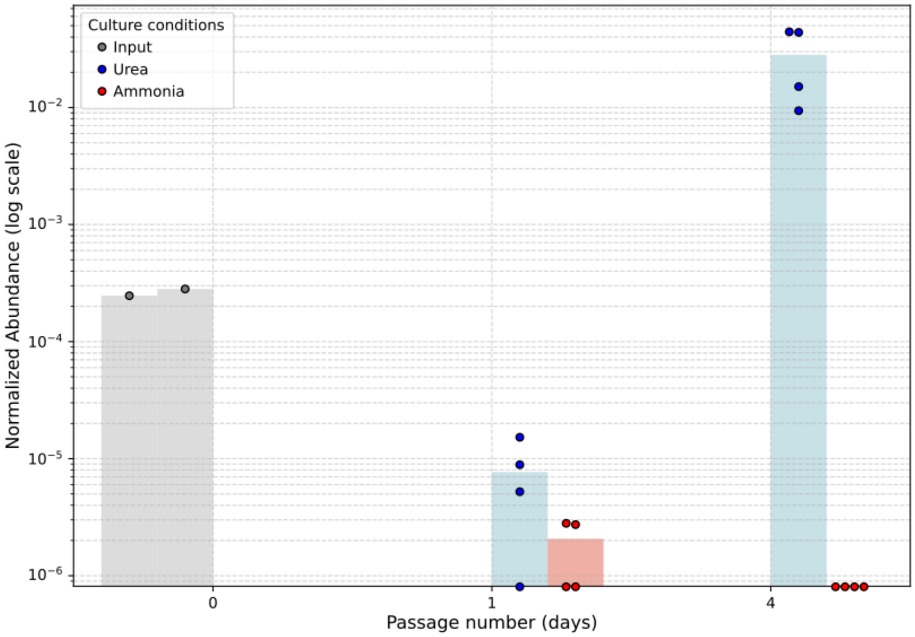


**Supplementary Figure 14. Transposon mutant frequency of a Glycine/betaine ABC transporter.**
Insertion site abundances of transposon mutants of the gene WP_115361527.1, annotated as a Glycine/betaine ABC transporter, in the transposon knockout library of *S. pasteurii* DSM 33 after growth on media supplemented with urea or ammonia.

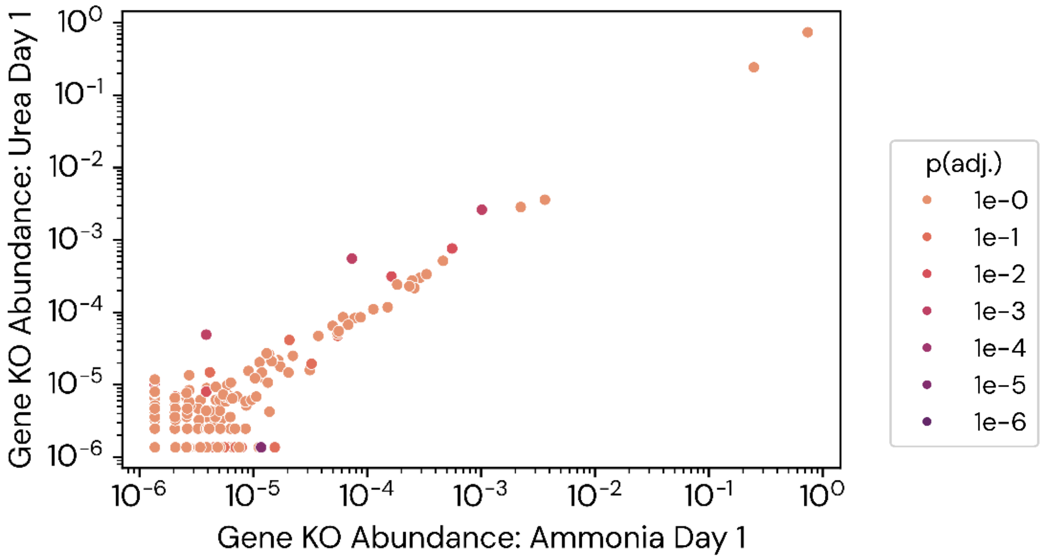


**Supplementary Figure 15. Abundances of transposon variants.**

Each point represents a gene. The relative abundance of insertions in that gene is plotted after a 24-hour passage of growth in media containing either urea or ammonia as an abundant nitrogen source.

### Supplementary Tables

**Supplementary Table 1. MicrobeMod analysis of *S. pasteurii* restriction-modification systems.**

Each row corresponds to a gene component of a restriction-modification (RM) system in *S. pasteurii* ATCC 11859 as identified by MicrobeMod[5]. The first column reports the RM system type of the corresponding operon, the second column reports the gene, the third column reports the gene family, the fourth column reports the associated homolog in the Restriction Enzyme Database (REBASE)[6], and the fifth column reports any observed methylation patterns in the genome associated with this RM system. Bolded bases indicate methylated base positions in the sequence motif.

| RM System | Gene | Gene family | REBASE homologue | Observed methylation |
|---|---|---|---|---|
| Type II R-M operon | DGCIAB_07790 | Methyltransferase | M.Spa337394ORF13930P | **G**AT**C**$_{m5}$ (92% of genomic sites methylated) |
| | DGCIAB_07795 | Restriction enzyme | Spa337394ORF13930P | **G**AT**C**$_{m5}$ (92% of genomic sites methylated) |
| Type I R-M operon | DGCIAB_15390 | Methyltransferase | M.Spa4822ORF1374P | Inactive |
| | DGCIAB_15395 | Specificity subunit | S.Spa4822ORF1374P | Inactive |
| | DGCIAB_15400 | Restriction enzyme | Spa4822ORF1374P | Inactive |

**Supplementary Table 2. Strains used in this study.**

| Strain | Genotype | Source | Growth medium |
|---|---|---|---|
| *Sporosarcina pasteurii* ATCC 11859 | WT | ATCC | $BNH_4YEM$ |
| *Sporosarcina pasteurii* DSM 33 | WT | DSMZ | $BNH_4YEM$ |
| *Sporosarcina* sp000813425 CVM220 | WT | This study | LB |
| *Escherichia coli* NEB10-beta | Δ(ara-leu) 7697 araD139 fhuA ΔlacX74 galK16 galE15 e14- ϕ80dlacZΔM15 recA1 relA1 endA1 nupG rpsL (StrR) rph spoT1 Δ(mrr-hsdRMS-mcrBC) | NEB (C3019) | LB |
| *Escherichia coli* BW29427 | thrB1004 pro thi rpsL hsdS lacZΔM15 RP4-1360 Δ(araBAD)567 ΔdapA1341::[erm pir] | B.L. Wanner* | LB + $DAP_{60}$ |
| *Escherichia coli* One Shot™ Pir1 | F–Δlac169 rpoS(Am) robA1 creC510 hsdR514 endA recA1 uidA(ΔMluI)::pir-116 | ThermoFisher Scientific (C101010) | LB |
| *Escherichia coli* One Shot™ Pir2 | F–Δlac169 rpoS(am) robA1 creC510 hsdR514 endA recA1 uidA(ΔMluI)::pir | ThermoFisher Scientific (C111110) | LB |
| *Escherichia coli* pir-116 | F - mcrA Δ(mrr-hsdRMS-mcrBC) ϕ80dlacZΔM15 ΔlacX74 recA1 endA1 araD139 Δ(ara, leu)7697 galU galK λ- rpsL (Str R) nupG pir-116(DHFR) | Biosearch Technologies (ECP09500) | LB |

*This strain is also available from the *E. coli* Genetic Research Center (14194)

**Supplementary Table 3. Summary of endogenous promoters used in this study.**

| Promoter ID | Gene ID | Usage in this study |
|---|---|---|
| P1 | Elongation factor Tu | ● Constitutive promoter characterisation<br>● Conversion to inducible promoter by tetO site introduction |
| P2 | 50S ribosomal protein L7/L12 | ● Constitutive promoter characterisation<br>● Expression of Himar/Tn5 transposase in Tn variant screen |
| P3 | Trigger factor | ● Constitutive promoter characterisation<br>● Conversion to inducible promoter by tetO site introduction |
| P4 | DNA-directed RNA polymerase subunit beta' | ● Constitutive promoter characterisation<br>● Expression of *tetR* for inducible promoter |
| P5 | 30S ribosomal protein S7 | ● Constitutive promoter characterisation |
| P6 | DNA-directed RNA polymerase subunit alpha | ● Constitutive promoter characterisation<br>● Expression of Himar/Tn5 transposase in Tn variant screen |
| P7 | 50S ribosomal protein L13 | ● Constitutive promoter characterisation<br>● Expression of Himar/Tn5 transposase in Tn variant screen |
| P8 | DNA polymerase III subunit beta | ● Constitutive promoter characterisation<br>● Expression of Himar/Tn5 transposase in Tn variant screen |
| P9 | Redox-regulated ATPase YchF | ● Constitutive promoter characterisation |
| P10 | Ribosome biogenesis GTPase Der | ● Constitutive promoter characterisation<br>● Expression of *tetR* for inducible promoter |
| P11 | L-serine ammonia-lyase, iron-sulfur-dependent subunit beta | ● Constitutive promoter characterisation<br>● Expression of Himar/Tn5 transposase in Tn variant screen |
| P12 | RelA/SpoT family protein | ● Constitutive promoter characterisation<br>● Expression of Himar/Tn5 transposase in Tn variant screen |
| P13 | LemA family protein | ● Constitutive promoter characterisation |
| P14 | DNA mismatch repair protein MutS | ● Constitutive promoter characterisation<br>● Expression of Himar/Tn5 transposase in Tn variant screen |
| P15 | Na/Pi cotransporter family protein | ● Constitutive promoter characterisation |
| P16 | GTP cyclohydrolase I FolE | ● Constitutive promoter characterisation |
| P17 | Division/cell wall cluster transcriptional repressor MraZ | ● Constitutive promoter characterisation |
| P18 | DNA primase | ● Constitutive promoter characterisation<br>● Expression of Himar/Tn5 transposase in Tn variant screen |
| P19 | NAD kinase | ● Constitutive promoter characterisation<br>● Expression of *tetR* for inducible promoter |
| P20 | Lipoate-protein ligase | ● Constitutive promoter characterisation |
| P21 | tRNA-Met | ● Constitutive promoter characterisation<br>● Conversion to inducible promoter by tetO site introduction |

**Supplementary Table 4. Promoter sequences used for design of inducible promoter libraries.**

Three endogenous promoters were selected for insertion of tetO sites and putative conserved promoter elements were identified: -35 box [consensus: TTGACA] in red, -10 box [consensus: TATAAT] in blue, and Shine-Dalgarno (SD) sequence [consensus: AGGAGG] in green. Motifs are annotated for each of the three promoter variants: P1, P3, and P21. Note that, as a tRNA-encoding promoter, P21 lacks an endogenous Shine-Dalgarno (ribosome-binding site) sequence.

| Promoter ID | Gene Identity | DNA Sequence |
|---|---|---|
| P1 | EF-Tu | TTTTCAGTATAAATAAGCTATAGAGTATAGGTGACTGGGGATTAATCTCCAGTACCTATAAAAATAAAAAACTTTCATATACTTAAGGAGGCTTTCCATA |
| P3 | Trigger factor | AATATATTGAAAGATCATGACGTTATGCTATACTGAAACAGTTGTCAAAATTGCATATACGTGTAAAATAGATTTACAAAAGATTCGGAGGTTTTTTAAC |
| P21 | tRNA-Methionine | CTATTAAAACTTTTCGACGCATTCCGAACTATTTTTCTATTTGCCGAAGTATTGACTTTATTTCAATAGATATGTAGTATAGTAAATGTTGCTTCAACAC |

**Supplementary Table 5. Transposase codon usage variants in transposon libraries.**
For details of codon-optimization and codon-harmonization processes, see **Methods**.

| Transposase | Unique ID | Codon usage type | Details |
|---|---|---|---|
| Himar | Himar.001 | Codon-optimized | Optimized based on *E. coli* codon usage pattern |
| | Himar.003 | Codon-harmonized | Harmonized using *E. coli* MG1655 codon usage pattern as source and *S. pasteurii* DSM 33 codon usage pattern as target |
| | Himar.004 | Codon-harmonized | Harmonized using *Stomoxys calcitrans* codon usage pattern as source and *S. pasteurii* DSM 33 codon usage pattern as target |
| | Himar.005 | Codon-harmonized | Harmonized using *Ectopseudomonas alcaliphila* IAM 14884 codon usage pattern as source and *S. pasteurii* DSM 33 codon usage pattern as target |
| | Himar.007 | Codon-optimized | Optimized based on *S. pasteurii* ATCC 11859 codon usage pattern |
| Tn5 | Tn5.001 | Codon-optimized | Optimized based on *E. coli* codon usage pattern |
| | Tn5.002 | Codon-optimized | Optimized based on *S. pasteurii* DSM 33 codon usage pattern |
| | Tn5.003 | Codon-harmonized | Harmonized using *E. coli* MG1655 codon usage pattern as source and *S. pasteurii* DSM 33 codon usage pattern as target |

### Supplementary Data Files

**Supplementary Data File 1 - Endogenous *Sporosarcina* promoters used in this study.**

**Supplementary Data File 2 - Design and sequences of inducible promoter library variants.**

**Supplementary Data File 3 - Details of all inducible and constitutive promoter library sequences included in oligo pool synthesis.**

**Supplementary Data File 4 - Plasmids used in this study.**

**Supplementary Data File 5 - Summary of sequence differences between *S. pasteurii* strains.**

**Supplementary Data File 6 - *Sporosarcina pasteurii* Tn-Seq experiment gene abundances.**

**Supplementary Data File 7 - Oligos used in this study.**

**Supplementary Data File 8 - Promoter induction statistics.**

All Supplementary Data Files are included as attachments under the ‘Supplementary material’ tab of the bioRxiv webpage for this manuscript.
